# Does Denoising Reduce Cross-Vendor Variability in Diffusion MRI? A Travelling Heads Evaluation

**DOI:** 10.64898/2026.11.24.768556

**Authors:** Francesco D’Antonio, Jose-Pedro Manzano-Patron, Olivier Mougin, Paul S. Morgan, Stamatios N. Sotiropoulos, Shaun Warrington

## Abstract

Between-scanner variability in magnetic resonance imaging (MRI)-derived features can be comparable to biological between-subject variability, even for the same individual scanned at different sites. MRI harmonisation is therefore an essential tool to ensure studies are generalisable across sites, reducing between-scanner variability and enabling pooling of data to achieve large sample sizes and more representative population samples. Denoising using MPPCA is widely used in preprocessing pipelines for diffusion MRI acquisitions to improve image SNR and it has been proposed as an implicit way of harmonising data. In this registered report, we hypothesised that denoising reduces cross-scanner variability and improves reliability in diffusion MRI metrics without changing cross-scanner bias, and that susceptibility-induced off-resonance reduces its efficacy. We tested these hypotheses using a travelling-heads resource of ten subjects scanned on six scanners from three vendors (GE, Philips, Siemens), with within-scanner repeats. We found that: i) Denoising reduced voxel-wise cross-scanner coefficients of variation for fractional anisotropy (FA), mean diffusivity (MD) and mean kurtosis (MK) (BF₁₀ > 100; median -3%, -9% and -32%). ROI-wise, variability decreased for MK (-12%), increased for FA (+8%), and evidence was inconclusive for MD. ii) Contrary to our hypothesis, denoising changed cross-scanner bias in every subject-scanner pair, by up to 3% of the mean; in exploratory analyses, this change was strongly related to contrast-to-noise ratio. iii) Evidence for a change in the registered measure of reliability was inconclusive, although exploratory analyses showed higher voxel-wise reliability after denoising. iv) There was no convincing evidence that off-resonance reduces denoising efficacy. Together, these results suggest that MPPCA denoising reduces the random, noise-driven component of cross-scanner variability, most strongly at the voxel level and for kurtosis, but does not consistently reduce systematic differences between scanners. As such, it should complement harmonisation, not replace it.

## 1. Introduction

Data harmonisation is a vital tool to ensure MRI-derived quantitative outcomes are generalisable across scanners and protocols, enabling multi-site studies and the pooling of data across studies (Bethlehem et al., 2022; Miller et al., 2016; Thompson et al., 2014; A. H. Zhu et al., 2025). This, in turn, enables larger and more diverse/representative samples. Large-scale MRI datasets are indispensable to assess population-wide trends in the brain, for example, exploring the effects in neurodevelopmental conditions (Jack et al., 2008) and brain ageing (Cole, 2020; Gaser et al., 2024; Mohammadi-Nejad et al., 2025), neuropsychiatric and mental health (van Erp et al., 2018), and for performing brain-wide association studies (Douaud et al., 2022). Brain effect sizes in such studies can be small but still biologically important; therefore, large sample sizes are often required to achieve sufficient statistical power and ensure generalisability (Button et al., 2013; Helmer et al., 2024; Marek et al., 2022; Marek & Laumann, 2025). However, to achieve large sample sizes and reach diverse populations, different sites and scanners are often used to acquire data. It is well established that cross-scanner acquisitions lead to differences in MRI measurements (Chen et al., 2014; Takao et al., 2011; Warrington et al., 2023, 2025; T. Zhu et al., 2011), necessitating the harmonisation of data to remove scanner-specific variations. Furthermore, as sample size increases, the sensitivity to confound effects also increases, potentially creating spurious correlations caused by confounds such as gradient heating or patient positioning (Alfaro-Almagro et al., 2021), further necessitating mitigation strategies.

Many harmonisation tools have been developed (Garcia-Dias et al., 2020; Johnson et al., 2007; Prevot et al., 2025; Radua et al., 2020), which seek to remove scanner-based variance in measurements whilst retaining biological variance (Radua et al., 2020; Tassi et al., 2024). However, depending on scanner heterogeneity, a large amount of data from each site is required to apply these tools (Parekh et al., 2022). Additionally, the effect of removing MRI-inherent thermal noise as a means to improve harmonisation has not been thoroughly explored.

In this study, we explored whether removing thermal noise in diffusion MRI (dMRI) acquisition using MPPCA denoising improves cross-scanner harmonisation, even when scanners are from different vendors. Furthermore, we evaluated the association of susceptibility artefacts with the ability of denoising to improve within-scanner harmonisation.

MPPCA is a patch-based denoising method for dMRI data (Veraart, Fieremans, et al., 2016; Veraart, Novikov, Christiaens, Ades-Aron, et al., 2016), commonplace in preprocessing pipelines (Ades-Aron et al., 2018; Chen et al., 2024; Huynh et al., 2024; Powell et al., 2022). At its core, this denoising method involves extracting 4-dimensional patches of diffusion-weighted data, rearranging them into a 2D matrix, and extracting signal-dominant principal components. To suppress noise-related components, a data-driven approach using the Marchenko-Pastur distribution (Marčenko & Pastur, 1967) is commonly used, where the threshold for signal-dominant eigenvalues is estimated from the distribution of a noise-only matrix of equivalent patch size (Veraart, Fieremans, et al., 2016; Veraart, Novikov, Christiaens, Ades-Aron, et al., 2016). Symmetric threshold estimation methods have been developed to more correctly estimate the noise-signal boundary (Cordero-Grande et al., 2019; Does et al., 2019; Olesen et al., 2023), and other methods use or estimate additional noise acquisitions to estimate the threshold (Henriques et al., 2023; Moeller et al., 2021). Recent work by Ades-Aron et al. (2025) shows that, by removing thermal noise using MPPCA, voxel-wise variation in dMRI-derived parametric maps decreases significantly across scanners from the same vendor. Therefore, in the context of large multi-site studies, MPPCA denoising could be a powerful tool to both harmonise data and improve image quality without increasing scan time (or reducing it without sacrificing image quality). However, the effectiveness of denoising in harmonisation across scanners of different vendors has yet to be tested.

Many hardware, image acquisition and reconstruction methods vary across scanner manufacturers (Gracien et al., 2020; Kruggel et al., 2010); therefore, it is important to assess whether the cross-scanner-within-vendor effectiveness of denoising is generalisable across vendors. For example, reconstruction of accelerated acquisitions (ARC/GRAPPA and ASSET/SENSE), gradient strength and slew rates, the number of head coil receiver channels, and bore diameters can vary significantly across vendors. Additionally, pulse sequence designs vary across manufacturers, influenced not only by hardware limitations but also by varying core pulse programming parameters, such as gradient raster timings (Fujita et al., 2025). The cumulative effect of these differences is highlighted in quantitative studies evaluating the variance of fundamental NMR measurements, such as T1 and T2 relaxation times, in standardised phantoms, showing variations across vendors (Boudreau et al., 2024), and variations in imaging-derived phenotypes (IDPs) measured from multi-vendor travelling head studies (Mirzaalian et al., 2016; Tax et al., 2019; Warrington et al., 2023, 2025). Vendor agnostic pulse sequence (Cordes et al., 2020; Karakuzu et al., 2022; Layton et al., 2017) and reconstruction software (Hansen & Sørensen, 2013; Uecker et al., 2016) have been developed to overcome vendor-specific sequence designs, enabling physics-based harmonisation (Fujita et al., 2025; Liu et al., 2024). However, much data has already been acquired and will continue to be acquired without the use of these tools. In addition, vendor-specific interpreters (used to convert vendor-agnostic sequences to vendor-specific ones) can introduce variance through sequence conversions and implementations, and some vendors are slow to make essential interpreters accessible to the research community (Roos et al., 2023). Finally, pulse sequence-based harmonisation in diffusion acquisitions would lead to using the lowest common value across scanners for important parameters, such as gradient strength and slew rate, leading to suboptimal acquisitions. Therefore, it is important to determine whether denoising can reduce cross-scanner, cross-vendor variance or whether removing noise will more clearly highlight the hardware- and software-related differences across manufacturers.

Additionally, previous work by Warrington et al. (2023) suggests that susceptibility-related artefacts could negatively affect the ability of denoising to reduce scan-rescan variability in dMRI. To avoid within-patch signal correlations and noise distribution modifications, optimal MPPCA denoising is performed prior to susceptibility correction. The work by Warrington et al. suggests that, in regions most severely affected by susceptibility artefacts and with large off-resonance fields, denoising did not reduce the variation of within-scanner repeats as much as in other brain regions. This is possibly due to signal shifts caused by susceptibility distortions, leading to spatially varying noise, breaking MPPCA’s statistical assumptions, and resulting in slightly different denoising and susceptibility corrections across repeats. Nevertheless, the evaluation in Warrington et al. involved region-of-interest (ROI) comparisons, in which the impact of denoising was significantly reduced, as regional averaging reduces noise in the first place (Ades-Aron et al., 2025). Furthermore, it is unclear if susceptibility-based detrimental effects are consistent across scanners.

Multiple travelling heads harmonisation resources have been developed and used to characterise scanner-related variability in multi-site diffusion MRI data (Ades-Aron et al., 2025; Cetin Karayumak et al., 2019; Grech-Sollars et al., 2015; Pohl et al., 2016; Tax et al., 2019; Warrington et al., 2023). These resources have been valuable to investigate cross-protocol, cross-scanner differences, including variability in hardware and acquisition parameters (Ades-Aron et al., 2025; Cetin Karayumak et al., 2019; Pohl et al., 2016; Tax et al., 2019). The ON-Harmony dataset (Warrington et al., 2023, 2025) provides a uniquely comprehensive resource, involving travelling heads scanned across scanners from three major vendors (GE, Philips, and Siemens), including within-scanner and between-scanner repeats of the same participants. The availability of within-scanner repeats provides a useful baseline against which to compare denoising results.

In this registered report we explored if denoising reduces cross-scanner-cross-vendor variability and improves reliability in dMRI metrics. We further explored whether susceptibility-induced off-resonance fields are affecting denoising efficacy. To test this, we used data from Phase B of the ON-Harmony dataset (Warrington et al., 2025) consisting of ten travelling heads acquisitions from six scanners across three vendors (GE, Philips, and Siemens) and containing six within scanner repeats from five scanners across the same three vendors.

## 2. Research Questions and Hypotheses

Using the ON-Harmony publicly available data (Warrington et al., 2025; https://doi.org/10.18112/openneuro.ds004712.v2.0.1), we tested the following pre-registered hypotheses (H1-H4 below). The accepted Stage 1 report is available at https://doi.org/10.17605/OSF.IO/MJ5KW.

**RQ1:** Does MPPCA denoising reduce cross-scanner variability in dMRI metrics?

**H1:** MPPCA denoising reduces subject-level variability in dMRI metrics when variability is computed across scanners from multiple vendors.

**a) Voxel-wise:** The voxel-wise subject-level coefficient of variation (CoV), computed across scanners for DTI and DKI metrics, will be reduced after MPPCA denoising.
**b) ROI-wise:** The ROI-wise subject-level CoV, computed across scanners for DTI and DKI metrics, will be reduced after MPPCA denoising.

**RQ2:** Does MPPCA denoising reduce cross-scanner bias in dMRI metrics?

**H2:** MPPCA does not affect subject-level cross-scanner bias in dMRI metrics.

**a) Voxel-wise:** The voxel-wise bias for DTI and DKI metrics will not be affected by MPPCA denoising.
**b) ROI-wise:** The ROI-wise bias for DTI and DKI metrics will not be affected by MPPCA denoising.

**RQ3:** Does MPPCA denoising improve the reliability of dMRI metrics across scanners?

**H3:** MPPCA improves the reliability of DTI and DKI metrics across scanners.

**a) Voxel-wise:** The voxel-wise reliability for DTI and DKI metrics will be increased after MPPCA denoising.
**b) ROI-wise:** The ROI-wise reliability for DTI and DKI metrics, will be increased after MPPCA denoising.

**RQ4:** Does susceptibility-related off-resonance moderate the effectiveness of MPPCA denoising in reducing within-scanner scan-rescan variability in dMRI-derived metrics?

**H4:** Higher susceptibility-related off-resonance frequencies are associated with a reduced effectiveness of MPPCA denoising in lowering within-scanner scan–rescan variability in DTI and DKI metrics.

**a) Voxel-wise:** The voxel-wise within-scanner subject-level CoV for DTI and DKI metrics after MPPCA denoising will show a positive association with mean off-resonance frequency, indicating reduced denoising-related CoV reduction in regions with higher off-resonance.
**b) ROI-wise:** The ROI-wise within-scanner subject-level CoV for DTI and DKI metrics after MPPCA denoising will show a positive association with mean off-resonance frequency, indicating reduced denoising-related CoV reduction in regions with higher off-resonance.

## 3. Methods

### 3.1 Data

We used Phase B of the openly available ON-Harmony dataset (https://openneuro.org/datasets/ds004712), which includes diffusion and anatomical (T1-weighted) MRI acquisitions from ten travelling heads scanned across six 3T scanners (Philips Ingenia, Philips Achieva, GE Premier 21ch head coil and GE Premier 48ch head coil, Siemens Prisma 32ch and Siemens Prisma 64ch head coil, with the two Prismas and Premiers being located at different sites). In addition, five of these participants had six within-scanner repeats, each subject using a different scanner (all scanners mentioned above, excluding the Philips Achieva). An overview of the scanners used is provided in Table 1, along dMRI acquisition protocols. For each scanning session, T1-weighted data were acquired using 1mm isotropic MPRAGE sequences.

**Table 1:** Overview of Scanner and Acquisition Parameters

| Scanner Name | Site | Bore Size (cm) | Maximum Gradient Strength (mT/m) | Gradient Slew Rate (T/m/s) | Head Coil Channels | Diffusion Weighted Sequence |
|---|---|---|---|---|---|---|
| Siemens Prisma | FMRIB, Oxford | 60 | 80 | 200 | 32 | 2mm iso, b= [0,1000,2000], 105 dirs, TE=92ms, TR=3.6s, PF=75% (zero-fill), PI=None, SMS=3 |
| Siemens Prisma | OHBA, Oxford | 60 | 80 | 200 | 64 | 2mm iso, b= [0,1000,2000], 105 dirs, TE=92ms, TR=3.6s, PF=75% (zero-fill), PI=None, SMS =3 |
| Philips Achieva | SPMIC-UP, Nottingham | 60 | Dual: 40 (80) | 200 (100) | 32 | 2mm iso, b= [0,1000,2000], 105 dirs, TE=70ms, TR=10.6s, PF=80% (homodyne-style), PI=2, SMS =None |
| Philips Ingenia | SPMIC-UP, Nottingham | 70 | 45 | 200 | 32 | 2mm iso, b= [0,1000,2000], 105 dirs, TE=98ms, TR=4.4s, PF=80% (homodyne-style), PI=1.5, SMS =3 |
| GE Premier | SPMIC-QMC, Nottingham | 70 | 80 | 200 | 48 | 2mm iso, b= [0,1000,2000], 105 dirs, TE=68ms, TR=4.9s, PF=80.8% (homodyne-style), PI=2, SMS =2 |
| GE Premier | OCMR, Oxford | 70 | 80 | 200 | 21 | 2mm iso, b= [0,1000,2000], 105 dirs, TE=70ms, TR=8s, PF=80.8% (homodyne-style), PI=2, SMS =2 |

DMRI data were acquired and reconstructed using vendor-specific pipelines according to each system’s implementation. Although the underlying reconstruction methods are well described in the literature, the exact vendor implementations are not fully disclosed. Parallel imaging (PI) was performed using SENSE-type methods for Philips, and ASSET for GE (no PI was used for Siemens acquisitions). Simultaneous multi-slice (SMS) acquisitions were implemented using multiband implementations for Siemens/Philips and HyperBand for GE. Partial Fourier (PF) data were reconstructed using vendor-specific approaches; homodyne-style reconstruction for Philips/GE and zero-filled reconstruction for Siemens, and apodisation filters were implemented in Philips and GE reconstructions. PF fraction for GE data is not available directly from the dicom header information and was calculated separately as shown in the Supplementary Material.

### 3.2 Researchers’ Prior Knowledge of the Data

At Stage 1, FD and JPMP had not observed the data. OM, PSM, SNS, and SW were involved in the initial acquisition and preprocessing of the raw data, as shown in Warrington et al. (2025), involving quality control checks, non-linear registrations to templates, and extraction of imaging-derived phenotypes for comparison across scanners. The preprocessing already carried out did not involve data generated from denoising or the comparison of susceptibility maps, and it involved non-linear registrations to standard templates (as opposed to linear registrations to subject-derived templates described below), meaning no key variables used in this analysis had been observed. Even though the raw data had been accessed and observed by some of the authors, none of the raw data had been denoised or pre-processed as described in the following section, meaning the data required for statistical analysis had yet to be generated, reducing the risk of bias. To further mitigate the risk of bias, conservative statistical thresholds were used to assess evidence for the hypotheses.

### 3.3 Data Preprocessing

For clarity and simplicity, the preprocessing steps common to testing all hypotheses are first described, followed by the specific additional processing steps for each hypothesis. An overview of the data processing is shown in Figure 1.

**Figure 1:**
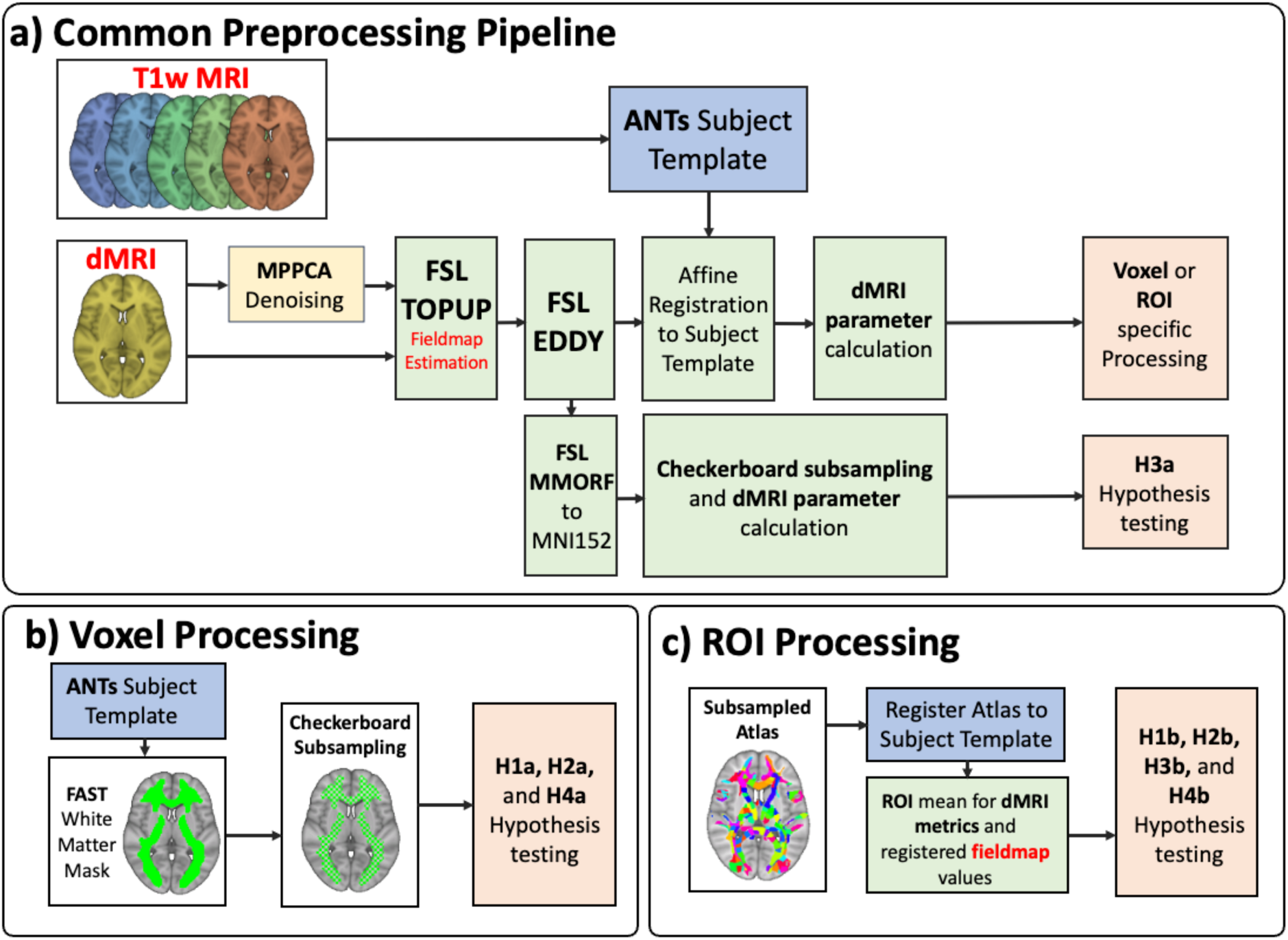
Preprocessing methods overview. **a)** Common preprocessing steps. DMRI images were susceptibility- and eddy-current-corrected, then linearly registered to a subject template. For H3a, eddy-corrected dMRI images were registered to MNI-152 space using MMORF. **b)** For voxel-wise analyses, a white matter mask was generated from the subject template, then subsampled using a checkerboard pattern to minimise cross-voxel correlations. ***c)*** *ROIs using the subsampled atlas were registered to the subject template using the inverse subject template to MNI152 transformation. Mean dMRI parametric values and susceptibility fieldmap values were calculated per ROI*.

## Common Preprocessing Pipeline

For all analyses, diffusion MRI data underwent susceptibility distortion correction (Andersson et al., 2003), eddy-current and motion correction (Andersson & Sotiropoulos, 2016), as part of an openly available preprocessing pipeline (Mohammadi-Nejad et al., 2020) using FSL v6.0.6.4 (this version was used throughout unless otherwise stated). Estimated motion per participant and across the cohort was estimated using FSL’s EddyQC (Bastiani et al., 2019), which is embedded in the preprocessing pipeline and reported in Supplementary Table S6. Deviation from Stage 1: EDDY was run using a GPU build from FSL 6.0.5. In one session, the reverse phase-encoded image was acquired left–right, so a custom susceptibility correction was used, identically in both conditions.

Two subject-specific anatomical templates were generated using T1-weighted scans. The first used all the subject’s sessions (used for H1, H2 and H3 processing), the second only used the six within-scanner repeats (used for H4). The subject-specific templates were created using ANTs (v2.4.4) multivariate template construction (Avants et al., 2008, 2011) (antsMultivariateTemplateConstruction2.sh). This approach used an initial linear registration to co-register the relevant T1-weighted scans, followed by iterative non-linear transformations to create an unbiased within-subject anatomical template.

Corrected dMRI datasets were linearly registered to their within-session T1-weighted anatomical acquisition using FSL’s FLIRT (Jenkinson et al., 2002) with six degrees of freedom, and subsequently linearly registered to the corresponding subject-specific template using the same method, with spline interpolation. To initially generate these transformations, they were carried out separately on the data. After being generated they were concatenated in a single transformation and applied to avoid resampling the data twice. Non-linear registrations were avoided to minimise blurring in the dMRI data. Fractional anisotropy (FA) and mean diffusivity (MD) maps were generated from the dMRI data using b=0 s/mm^2^ and b=1000s/mm2 diffusion shells and mean kurtosis (MK) using all b-value shells in subject template space. Deviation from Stage 1: Stage 1 did not specify exclusion criteria for metric values. To retain physically plausible values, we kept, separately for each metric, only voxels with FA > 0, MD > 0 or 0 < MK ≤ 3 respectively (Jensen & Helpern, 2010; Tabesh et al., 2011), in both the denoised and non-denoised data and in all sessions relevant to each comparison. Excluded voxels were also left out of ROI means. Supplementary Table S7 gives the units kept.

An independent copy of the dMRI data was denoised with MPPCA prior to preprocessing, yielding parallel denoised FA, MD and MK maps in subject template space. MPPCA denoising was carried out using eigenvalue thresholds defined in Cordero-Grande et al. (2019), optimal singular value shrinkage using the Frobenius norm (Gavish & Donoho, 2017), variable patch sizes based on local signal similarity (Ades-Aron et al., 2025), as implemented in the DESIGNER pipeline (v2.0.11) (Ades-Aron et al., 2018; Chen et al., 2024). A table of scripts with optional parameters which were registered for use at Stage 1 can be found in the Supplementary Materials, and the full preprocessing pipeline can be found in the project repository (https://github.com/SPMIC-UoN/harmonisation-with-denoising). As the parametric maps were derived from a single-session acquisition, no B1 field inhomogeneity correction was carried out as the fractional parametric maps are robust to static multiplicative inhomogeneities.

## H1a: Voxel-wise, Cross-Scanner Variability

For each of the ten travelling heads scanned across the six scanners, voxel-wise coefficients of variance (CoV) were calculated for FA, MD and MK maps, separately for denoised and non-denoised data:

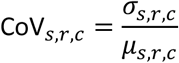

Where *μ_s_*_,*r*,*c*_ and *σ_s_*_,*r*,*c*_ are the mean and standard deviation of FA, MD or MK values for subject *s*, voxel/ROI *r*, and denoising condition *c* (denoised or non-denoised), across scanners. In the case where a subject has multiple repeats for a scanner, a mean of the repeats was calculated.

A white matter mask was generated from the anatomical template using FSL’s FAST (Zhang et al., 2001), using a white matter partial volume threshold of 0.5, and subsampled with a checkerboard pattern to reduce voxel dependence for FA, MD and MK parametric maps. Deviation from Stage 1: To match the sample size of the sparse checkerboard used in the data-adequacy simulations (∼6,800 voxels; Section 3.6), one white matter voxel in eight was kept (alternate voxels in each dimension; 6,301–9,397 per subject before exclusions). Of 36 voxel-wise verdicts, 35 were the same with a denser checkerboard (∼50% of voxels; Supplementary Table S18); the exception was H4a for MD (Ingenia32), which supported H4 only with the denser checkerboard.

## H1b: ROI-Wise, Cross-Scanner Variability

ROI based analyses were carried out by parcellating the 42 distinct tract ROIs obtained from the XTRACT WM atlas (Warrington et al., 2020), available in FSL. This atlas is selected for its broad brain coverage, including the anterior and inferior regions of the brain, which are more prone to susceptibility artefacts.

Atlas parcellation was carried out to increase homogeneity in ROI size and improve localisation. The subsampling iteratively splits the original tract labels by identifying principal directions of the tracts and merging small splits into adjacent larger ROIs, while still maintaining the original cross-tract boundaries, to achieve a mean ROI size of 400 voxels, with a threshold for the smallest acceptable ROI size of 200 voxels. The code used to parcellate the original atlas and the resulting parcellated atlas are available in the GitHub repository (https://github.com/SPMIC-UoN/harmonisation-with-denoising).

Non-linear registration warp fields were derived between each of the subjects’ T1w templates and the standard space MNI152 T1w brain using FSL’s FNIRT (Andersson, 2010). These were subsequently used to bring the subsampled ROIs into the subject template spaces. Mean FA, MD and MK parameters were calculated within each ROI, thresholding for a value of 0.5 per ROI. Furthermore, voxels outside the subject-derived white matter mask were not considered. CoV across scans (for each ROI) were calculated for denoised and non-denoised versions of the data.

## H2: Cross-Scanner Bias

To assess bias, we used a combination of the within-scanner and cross-scanner acquisitions. Since each set of scanner repeats involved a different subject, the cross-scanner bias was confounded with subject differences. Therefore, changes in bias were calculated separately for each of the five scanners-subject pairs. Following the common preprocessing pipeline, mean dMRI parametric maps were generated from within-scanner repeats after registration to subject templates (*μ_r_*_,c_^WITHIN^) and from acquisitions across scanners (*μ_r_*_,c_^ACROSS^, not including the within-scanner repeats used to calculate *μ_r_*_,c_^WITHIN^) for each subject with scan-rescan acquisitions for voxel/ROI *r* and denoising condition *c*. The bias for a subject/scanner pair was calculated by taking the difference of these values (Bias*_r_*_,*c*_), and a change in bias resulting from the denoising condition can then be evaluated (Δ*_r_*^Bias^):

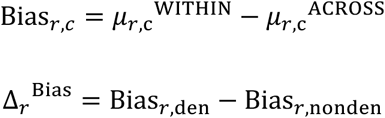

Like the processing described in H1, a white matter mask was subsampled for the voxel-wise analysis and ROIs registered for the ROI-wise analysis using within subject templates.

## H3: Cross-Scanner Reliability

For the voxel-wise analysis of the effect of denoising on cross-scanner reliability of dMRI metrics, all dMRI sessions were registered to a common space using non-linear transformations to go from subject-native to MNI152 template space. To do so whilst minimising blurring, multi-modal registration from subject space to MNI152 space was carried out using FSL’s MMORF (Lange et al., 2024), using FA and DTI derived tensors alongside the T1w image. For the ROI-wise analysis, the ROI labels provide cross-subject correspondence, meaning only within subject registration was required (instead of registering to MNI152 space).

## H4: Denoising Association with Off-Resonance Fieldmaps

In addition to the common preprocessing steps, voxel-wise off-resonance frequency maps were generated during susceptibility correction using FSL TOPUP for each of the five scanners with six single-subject within-scanner repeats. These fieldmaps were linearly registered to the subject anatomical template using the same affine transformations applied to the corresponding dMRI data. Voxel/ROI-wise CoV values were calculated using single-subject repeats for each scanner separately to avoid scanner/subject confounds as described in H2.

Both denoised and non-denoised pipelines produced registered fieldmaps in subject template spaces. Mean off-resonance frequency values were computed per voxel and used in subsequent analyses alongside FA, MD and MK CoV measures.

### 3.4 Statistical Analysis

#### H1: Cross-Scanner Variability

To test whether MPPCA denoising reduces variability across scanners, we fitted a mixed-effects Bayesian model with random intercepts for subjects and fixed effects for denoising condition to voxel-(**H1a**) or ROI-wise (**H1b** – using ROIs as an additional random effect) coefficients of variance (CoV) for FA, MD and MK. The model was run independently for each parametric map. Prior to modelling, parametric CoV values were z-standardised across subjects and voxels or ROIs. Each observation *y_s_*_,*r*,*c*_ corresponds to a (standardised) CoV value for a given subject (*s*), voxel (*r*), and condition (*c*: non-denoised or denoised). The model estimated a population-level denoising effect *α*_1_ while accounting for between-subject differences:

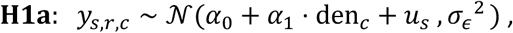

where *α*_0_ is the population mean CoV, *u_s_* ∼ N(0, *σ_u_*^2^) are random intercepts for subjects, den*_c_* ∈ {0,1} is a binary indicator of denoising condition (0 = non-denoised, 1 = denoised), and *σ_ε_* is the residual standard deviation capturing within-subject variability not explained by the model. Since the goal of the analysis was to understand if there is a population averaged denoising effect, not the variability across subjects, and given the relatively low number of subjects available for the analysis, random slopes allowing the denoising effect to vary by subject were not included.

For the ROI based comparison, an additional term is added to model random intercepts for ROIs, *v_r_* ∼ N(0, *σ_v_*^2^).

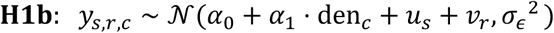

Evidence for an effect of denoising was assessed using Bayes factor comparing models with and without the denoising term.

## H2: Cross-Scanner Bias

To assess the effect of denoising on between scanner bias, the following model was fitted to each scanner/subject pair for the voxel-wise (H2a) and ROI-wise (H2b) analysis:

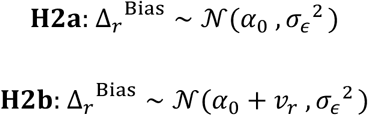

For H2, evidence for an effect of denoising on cross-scanner bias was assessed using Bayes factors comparing a model including the intercept parameter *α*_0_to a point-null model where *α*_0_ = 0.

## H3: Cross-Scanner Reliability

To assess reliability, the following mixed effects model was fitted for denoised and non-denoised data:

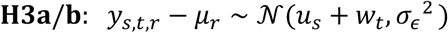

Where *μ_r_* is the mean value across all sessions for voxel/ROI *r*, *u_s_* ∼ N(0, *σ_sub_*_j*ects*_^2^) and *w_t_* ∼ N(0, *σ_scanners_*^2^) are the variance between subjects and variance between scanners respectively. Reliability was assessed via the intraclass correlation coefficient (ICC):

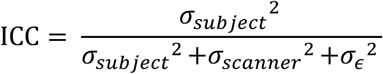

Posterior samples of variance components are obtained via Bayesian inference, yielding posterior distributions for each component. ICC was computed for each posterior draw, resulting in a posterior distribution of ICC for each condition. The difference in reliability due to denoising condition is then defined as:

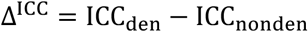

Which was calculated for each posterior draw, creating a posterior distribution for Δ^ICC^. The effect of denoising was assessed using the posterior distribution of Δ^ICC^.

## H4: Denoising Association with Off-Resonance Fieldmaps

To test the interaction between off-resonance frequency and denoising we fitted a Bayesian regression model to the voxel-wise (**H4a**) or to the ROI-wise (**H4b**) coefficients of variance (CoV) for FA/MD/MK with the corresponding off-resonance frequency fieldmap values. Since each set of scanner repeats involves a different subject, modelling the cross-scanner differences was confounded with subject differences. Therefore, the model was fitted separately for each of the five scanner-subject pairs. Additionally, since denoising occurs prior to susceptibility and eddy-current corrections, the estimated corrections could themselves change because of denoising. To evaluate if these effects should be taken into consideration within the model, a separate independent dataset (not involved in the future analysis) of six within-scanner repeats from Phase A of the ON-Harmony data was taken using the Siemens Trio (a scanner not evaluated in the future analysis) to compare off-resonance frequency maps and eddy current corrections metrics between denoised and non-denoised scans. Detailed results are found in supplementary material and show that the correction maps do not significantly vary with denoising condition. Therefore, differences in eddy current correction or susceptibility correction contributions across non-denoised/denoised conditions were not considered in the model. As there is no detectable difference between non-denoised/denoised off-resonance frequency maps, a mean of the two was calculated per session.

To test whether the effectiveness of MPPCA denoising depends on off-resonance field strength, we computed the difference in standardised CoV between the denoised and non-denoised parametric maps for each voxel/ROI (*r*):

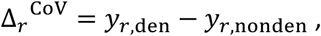

where *y_r_*_,*c*_ denotes the z-standardised CoV value for voxel/ROI *r* under condition *c* (non-denoised or denoised). Each scanner was analysed separately using a Bayesian regression model:

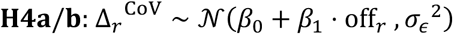

where off*_r_* denotes the mean voxel/ROI off resonance fieldmap value across repeats, *β*_0_ represents the mean denoising effect when the off-resonance field is zero (i.e., the expected change in standardised CoV attributable to denoising at zero off-resonance), and *β*_1_ quantifies how off-resonance field strength modulates the magnitude of denoising effect. Deviation from Stage 1: Stage 1 did not specify whether the magnitude or the signed value of off-resonance would be used. Because H4 concerns whether the strength, not the sign, of the off-resonance affects denoising, off*_r_* was calculated by taking the magnitude of the off-resonance (Hz) in each repeat and condition and then averaging these magnitudes (and, ROI-wise, averaging across voxels). This choice was made before any model was fitted.

## Priors and Inference Framework

Weakly informative priors were used to regularise estimates. Fixed effects (e.g., *α*_0_, *α*_1_, *β*_0_, *β*_1_) are assigned N(0,1) priors. Standard deviations of random effects and residuals (e.g., *σ_sub_*_j*ect*_, *σ_scanner_*, *σ_ε_*) are assigned Half-Normal(0,1) priors. Posterior distributions were estimated using Hamiltonian Monte Carlo sampling (4 chains, 4000 iterations, warmup = 2000). We report posterior means, 95% credible intervals, and Bayes factors comparing models with and without the relevant terms when BF is used to evaluate the hypothesis. Convergence was assessed using *R*Q < 1.01 and effective sample size > 1000.

### 3.5 Result Interpretation

#### H1: Cross-Scanner Variability

We interpreted the evidence using Bayes factors with BF10 > 10 and posterior distribution means of *α*_1_ being negative as strong evidence for the respective hypotheses, calculated for FA/MD/MK and voxels/ROIs separately. BF10 < 1/10 was interpreted as strong evidence for the null hypotheses, indicating that thermal noise does not significantly contribute to cross-scanner variability, suggesting hardware and software differences, not addressed by denoising, contribute more strongly to cross-scanner variability.

## H2: Cross-Scanner Bias

Evidence for no change in bias was assessed via BF10. BF10 < 1/10 was interpreted as strong evidence for the null hypothesis (no systematic bias introduced by denoising). BF10 > 10 was interpreted as evidence that MPPCA introduces systematic bias into parametric maps.

## H3: Cross-Scanner Reliability

Evidence for increased reliability following denoising was assessed via the posterior probability *P*(Δ^ICC^ > 0), with values greater than 0.95 indicating support for H3 (increase in reliability), values below 0.05 indicating support for the null (reduction of reliability), and intermediate values considered inconclusive.

## H4: Denoising Association with Off-Resonance Fieldmaps

We interpreted the evidence using Bayes factors with BF10 > 10 and posterior distribution means of *β*_1_ being positive, as strong evidence for the respective hypotheses, calculated for FA/MD/MK and voxels/ROIs separately. Results for each scanner are reported separately and interpretations are carried out for each scanner/subject couple independently, as subject and scanner/vendor differences are confounded. BF10 < 1/10 are interpreted as strong evidence for the null hypothesis, suggesting denoising’s ability to reduce within-scanner variability is not moderated by susceptibility artefacts. BF10 values between the specified ranges (10 > BF10 > 1/10) are interpreted as inconclusive evidence.

For all hypotheses, within hypotheses differences driven by voxel or ROI analysis could occur. For example, in H1 results could show BF10 > 10 in voxel-wise analysis (H1a) but below the significance threshold in ROI based analysis (H1b), or with a lower mean posterior distribution. This would support previous evidence that by taking ROI averages, we are already averaging-out thermal-noise random variance, making MPPCA preprocessing less effective in harmonisation. Similarly for other hypotheses, differences driven by voxel or ROI analysis were interpreted as taking ROI averages are averaging-out thermal-noise random variance, reducing the efficacy of denoising.

### 3.6 Bayesian Data Adequacy Evaluation

To give probabilistic assurance that we had adequate data to detect an effect of interest, each of the three models was simulated using 4 chains, 4000 iterations, warmup = 2000. To estimate an exemplary sample size in voxel-wise simulations, a sparse white matter checkerboard mask was generated from the MNI152 template using the same processing steps outlined in Section 3.3, obtaining a sample size of 6800 voxels. Expected effect size for H1 was taken as 0.1, for H3 going from *σ_ε_* = 0.6 → 0.3 due to denoising and 0.2 for H4, as conservatives estimate from previous work (Ades-Aron et al., 2025; Warrington et al., 2023) and a sizable effect size of interest. Cross-subject baseline variability was set to *u_s_* = 0.6 based on empirical cross-subject standard deviation of z-scored quality metrics from Phase A of the ON-Harmony dataset, using values from dMRI reflecting noise which could affect denoising (Avg. SNR, Avg. CNR, Avg SNR Std, Avg CNR Std, absolute motion, and relative motion). Between scanner variability was taken as slightly lower than between subject variability (*w_t_* = 0.4) from previous evidence (Warrington et al., 2023). ROI-level random intercepts were assigned smaller standard deviations (*v_r_* = 0.2) than subject-level effects to reflect secondary variability due to ROI size and location relative to dominant between-subject heterogeneity. The code used for these simulations and full results from the simulations can be found in the GitHub repository (https://github.com/SPMIC-UoN/harmonisation-with-denoising). Table 2 shows the fraction of simulated repetitions with statistical tests satisfying the significance thresholds for each model and their respective parameters used. The simulations showed a probability of detecting effect sizes of interest greater than 0.8 for the sample sizes and effect sizes outlined above, giving probabilistic assurance that we have adequate data to detect an effect of interest.

**Table 2:**
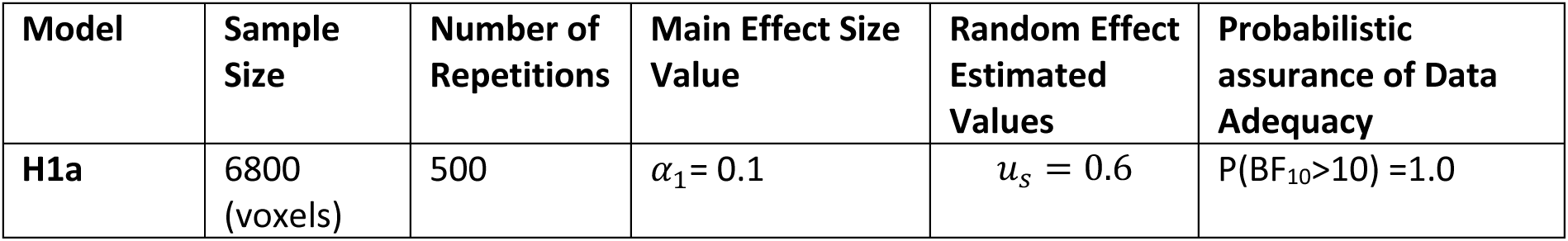
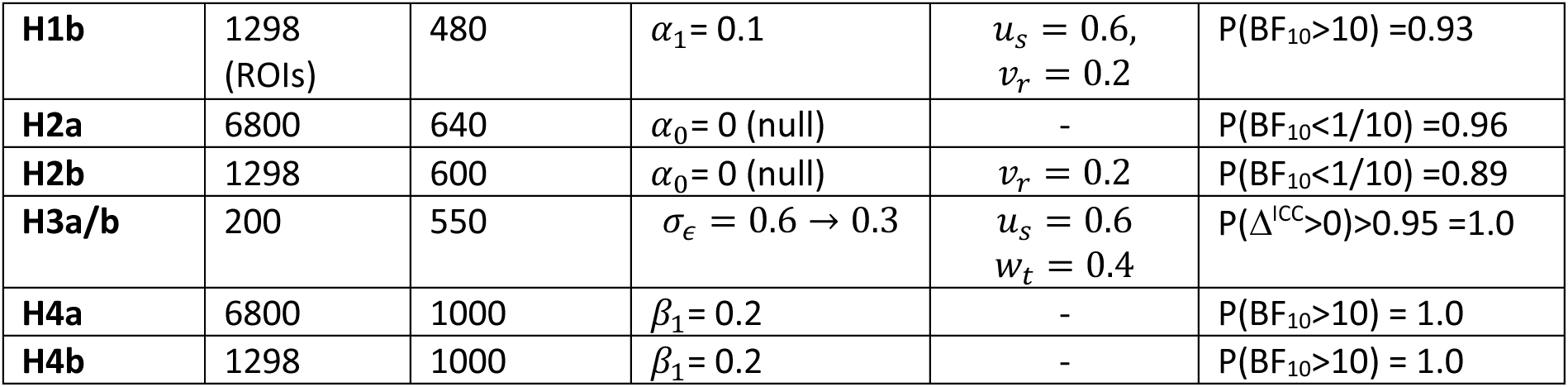
Simulated Results for Data Adequacy Evaluation

| **Model** | **Sample Size** | **Number of Repetitions** | **Main Effect Size Value** | **Random Effect Estimated Values** | **Probabilistic assurance of Data Adequacy** |
|---|---|---|---|---|---|
| **H1a** | 6800<br>(voxels) | 500 | *α*<sub>1</sub>= 0.1 | *u*<sub>*s*</sub> = 0.6 | P(BF<sub>10</sub>>10) =1.0 |
| **H1b** | 1298<br>(ROIs) | 480 | *α*<sub>1</sub>= 0.1 | *u*<sub>*s*</sub> = 0.6,<br>*v*<sub>*r*</sub> = 0.2 | P(BF<sub>10</sub>>10) =0.93 |
| **H2a** | 6800 | 640 | *α*<sub>0</sub>= 0 (null) | - | P(BF<sub>10</sub><1/10) =0.96 |
| **H2b** | 1298 | 600 | *α*<sub>0</sub>= 0 (null) | *v*<sub>*r*</sub> = 0.2 | P(BF<sub>10</sub><1/10) =0.89 |
| **H3a/b** | 200 | 550 | *σ*<sub>*ϵ*</sub> = 0.6 → 0.3 | *u*<sub>*s*</sub> = 0.6<br>*w*<sub>*t*</sub> = 0.4 | P(Δ<sup>ICC</sup>>0)>0.95 =1.0 |
| **H4a** | 6800 | 1000 | *β*<sub>1</sub>= 0.2 | - | P(BF<sub>10</sub>>10) = 1.0 |
| **H4b** | 1298 | 1000 | *β*<sub>1</sub>= 0.2 | - | P(BF<sub>10</sub>>10) = 1.0 |

## 4. Results

### 4.1 Registered Analysis

#### Data and Modelling Quality Control

Denoising did not meaningfully change the susceptibility or eddy-current corrections (Supplementary Figures S8-9). EDDY’s estimated RMS motion was slightly lower with denoising (−0.049mm) but far below the 2mm voxel size. Motion estimated by EDDY in the non-denoised data was lower than in the UK Biobank (absolute motion 0.70 ± 0.54 mm vs 1.09 ± 0.47 mm; UK Biobank Showcase Data-Field 24450, processed as in Alfaro-Almagro et al., 2018).

Voxels with unphysical metric values were excluded (Section 3.3; Supplementary Table S7). Voxel-wise samples averaged 7,230–7,841 voxels per subject (H1a) and 7,303–8,294 per subject–scanner pair (H2a, H4a), above the 6,800 assumed in the data-adequacy simulations; H3a used 9,163 (FA), 9,031 (MD) and 2,668 (MK) voxels. ROI-wise samples (1,143–1,285 ROIs per subject, subject–scanner pair or H3 model) were slightly below the 1,298 assumed. Sixty-nine of 78 fits met the convergence criteria for the parameters used for inference. The other nine (seven H2b fits and two H3 fits, affecting two H3 tests) are reported as fitted and marked †.

## H1: Cross-Scanner Variability

Results are shown in Figure 2 and Supplementary Table S10. Voxel-wise, denoising reduced cross-scanner CoV for all three metrics (Figure 2, top row; FA −0.036 [−0.046, −0.026]; MD −0.074 [−0.084, −0.064]; MK −0.425 [−0.436, −0.415]; BF₁₀ > 100 for all). In native units, the median voxel-wise CoV decreased from 13.0% to 12.6% for FA, from 6.4% to 5.8% for MD, and from 10.0% to 6.8% for MK, and CoV was lower after denoising in 56%, 70%, and 81% of voxels respectively. The effects for FA and MD were smaller than the effect of interest used in the data-adequacy simulations (0.1).

**Figure 2.**
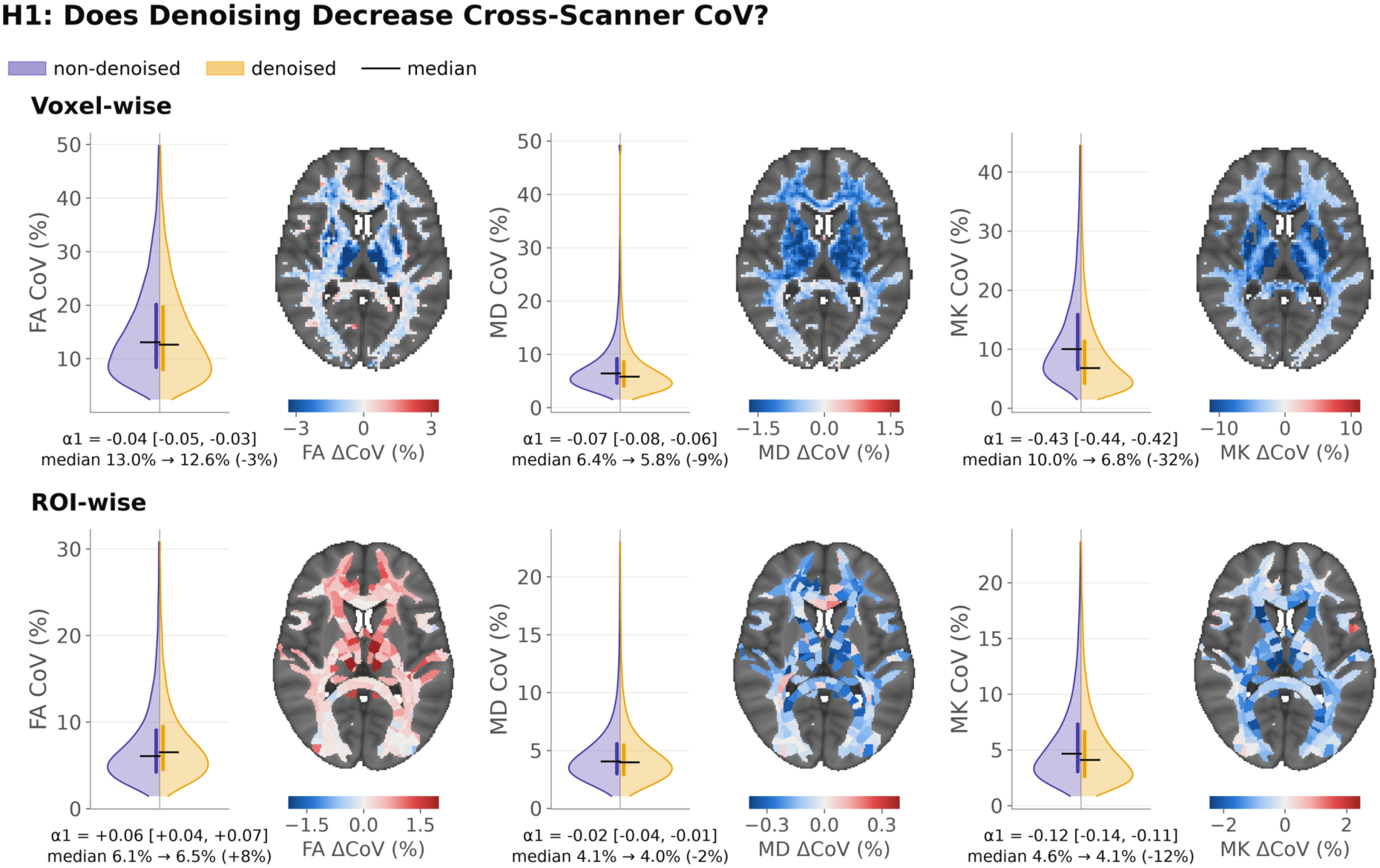
H1: Cross-scanner variability with and without MPPCA denoising. Coefficient of variation (CoV) of each unit (voxels/ROIs) across the six scanners. Non-denoised (left half, violet) and denoised (right half, yellow). Top row voxel-wise (white-matter checkerboard voxels), bottom row ROI-wise (ROI means). Black line = median, bars = interquartile range. For display, the voxel-wise map shows all white-matter voxels; the analysis used only checkerboard-subsampled voxels. ΔCoV = denoised − non-denoised (blue = lower with denoising).

ROI-wise (Figure 2, bottom row), there was strong evidence for H1b for MK (α1 = −0.123 [−0.141, −0.106], BF₁₀ > 100; median CoV 4.6% to 4.1%). Evidence for MD was inconclusive (α1 = −0.023 [−0.040, −0.005], BF₁₀ = 0.23; median 4.1% to 4.0%). For FA, there was evidence of an effect in the direction opposite to H1b (α1 = +0.060 [+0.044, +0.075], BF₁₀ >100): median ROI-wise FA CoV increased from 6.1% to 6.5%. The increase in FA ROI CoV is analysed in the exploratory analyses.

## H2: Cross-Scanner Bias

Contrary to H2, there was strong evidence that denoising changed cross-scanner bias (Figure 3, Supplementary Table S11). The change was large relative to the variability of ΔBias across units (|α0| = 0.10-0.51 voxel-wise and 0.36-1.23 ROI-wise) but small in native units: the mean ΔBias was between -3.0% and +2.4% of the mean value for FA, -0.4% and +0.4% for MD, and -1.7% and +1.5% for MK. The direction depended on the repeat scanner and was matched in direction at the voxel and ROI levels. Seven ROI-wise fits did not meet the convergence criteria (†); all had BF₁₀ > 100. The relationship between change in bias and scanner CNR was assessed in exploratory analyses.

**Figure 3.**
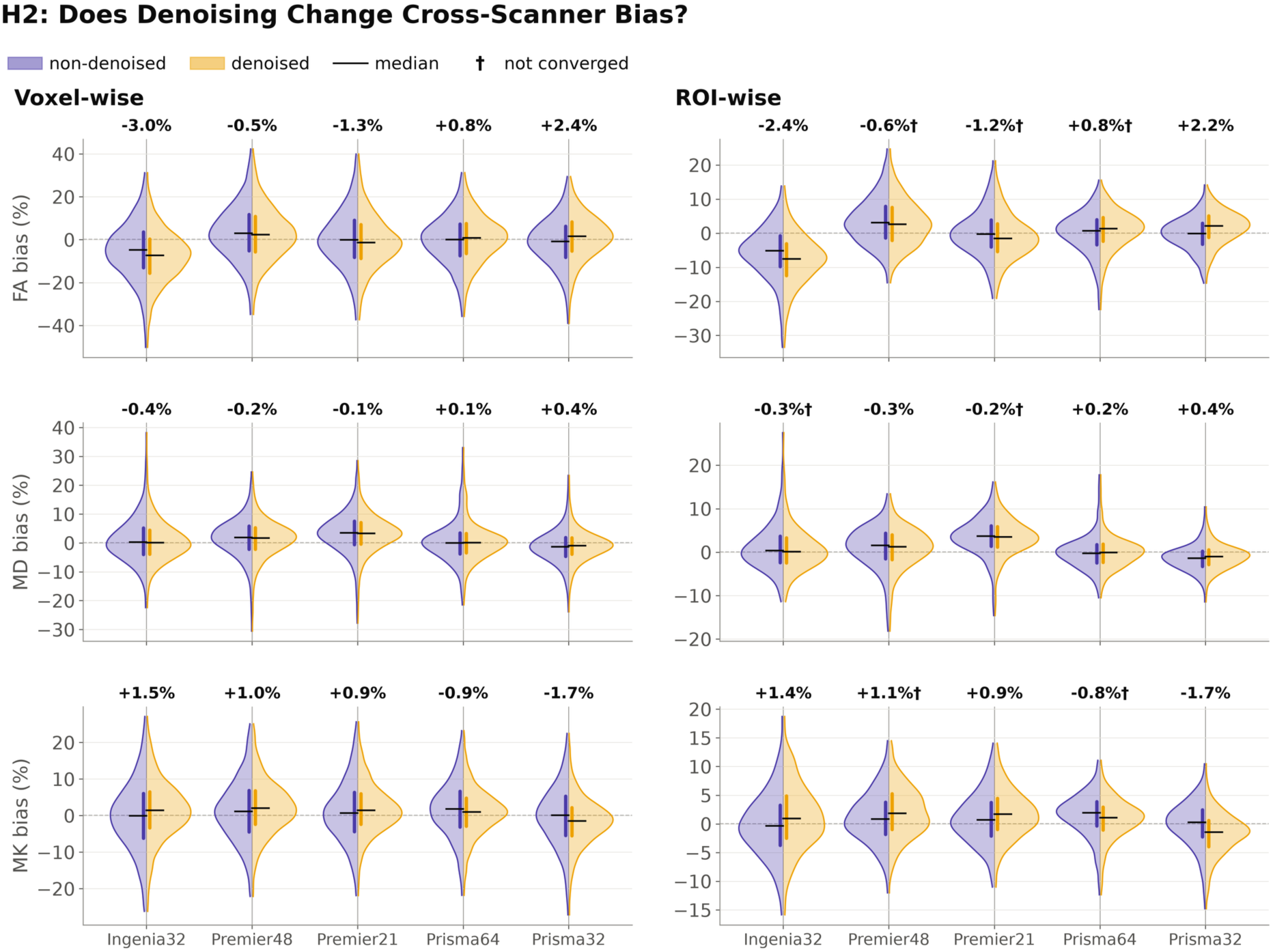
H2: Cross-scanner bias with and without MPPCA denoising. Five repeat pairs (one subject scanned six times on one scanner). Per-unit bias = mean over the six repeat sessions − mean over the same subject’s other five scanners. Non-denoised (left half, violet) and denoised (right half, yellow). Black line=median, bar=interquartile range. Numbers above each pair: ΔBias = denoised − non-denoised bias (% of mean). † did not meet the convergence criteria.

## H3: Cross-Scanner Reliability

The pre-registered ICC was low in both arms (Figure 4, Supplementary Table S12), indicating that, with one subject and one scanner effect shared across all voxels/ROI, most of the variance was attributed to the residual component. Posterior mean ΔICC was close to zero or positive for all six tests, and the posterior probability of an increase ranged from 0.50 to 0.81, below the 0.95 threshold. Evidence was therefore inconclusive for H3a and H3b for all metrics.

**Figure 4.**
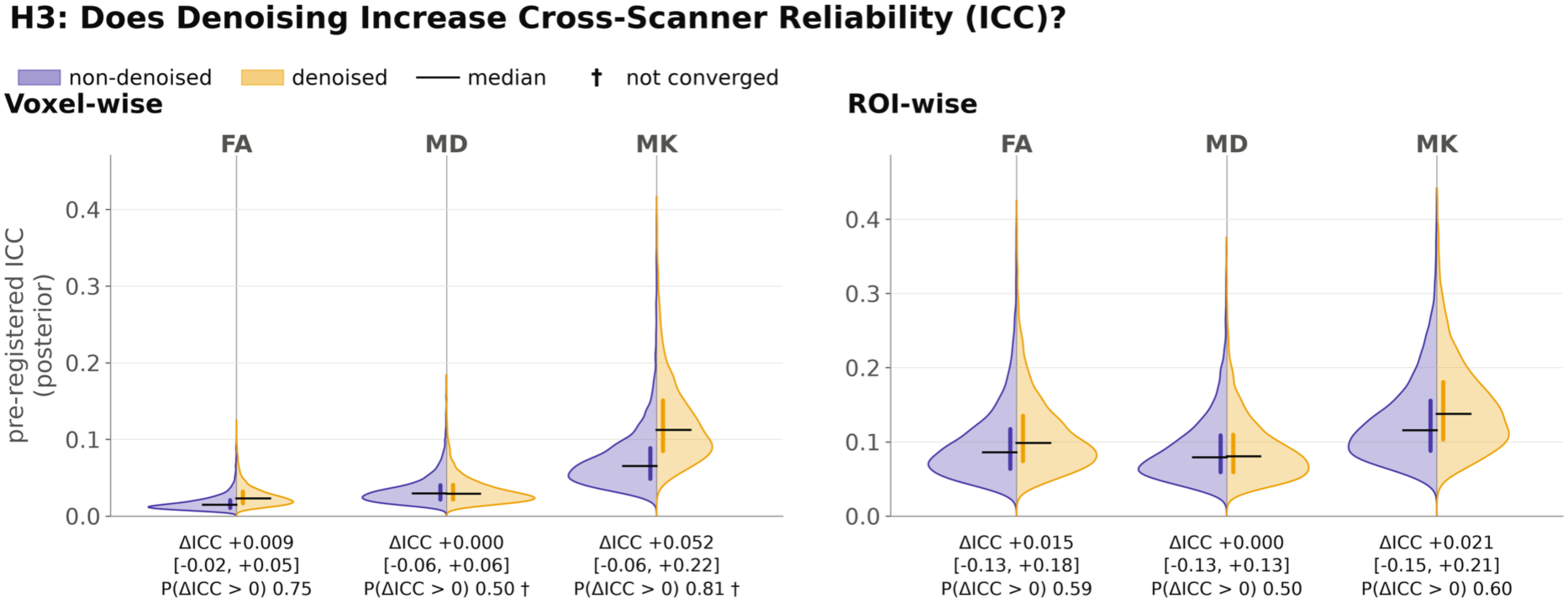
H3: Cross-scanner reliability with and without MPPCA denoising. Posterior distribution of the pre-registered ICC, fitted separately to non-denoised (left half, violet) and denoised (right half, yellow) data. † did not meet the convergence criteria.

## H4: Denoising Association with Off-Resonance Fieldmaps

Evidence for moderation was mixed across the tests (Figure 5; Supplementary Table S13). Five supported H4, 11 supported the null, and 14 were inconclusive. A visualisation of the mean off-resonance fields and change in CoV for MK is shown in Figure 5A. Twelve of the inconclusive tests showed evidence in the opposite direction (β1 < 0, BF₁₀ > 10). Voxel-wise, β1 (per SD of |off-resonance|) was at most 30% of the denoising effect β0 in magnitude (Figure 5B, C). ROI-wise, it was comparable to or larger than β0 for some pairs, particularly where β0 was close to zero (Figure 5B). The data-adequacy simulations assumed standardised off-resonance and β1 = 0.2; all |β1| per SD were smaller (at most 0.16). In the exploratory analysis, we assessed how large an association the spatial smoothness of the off-resonance field could produce by chance.

**Figure 5.**
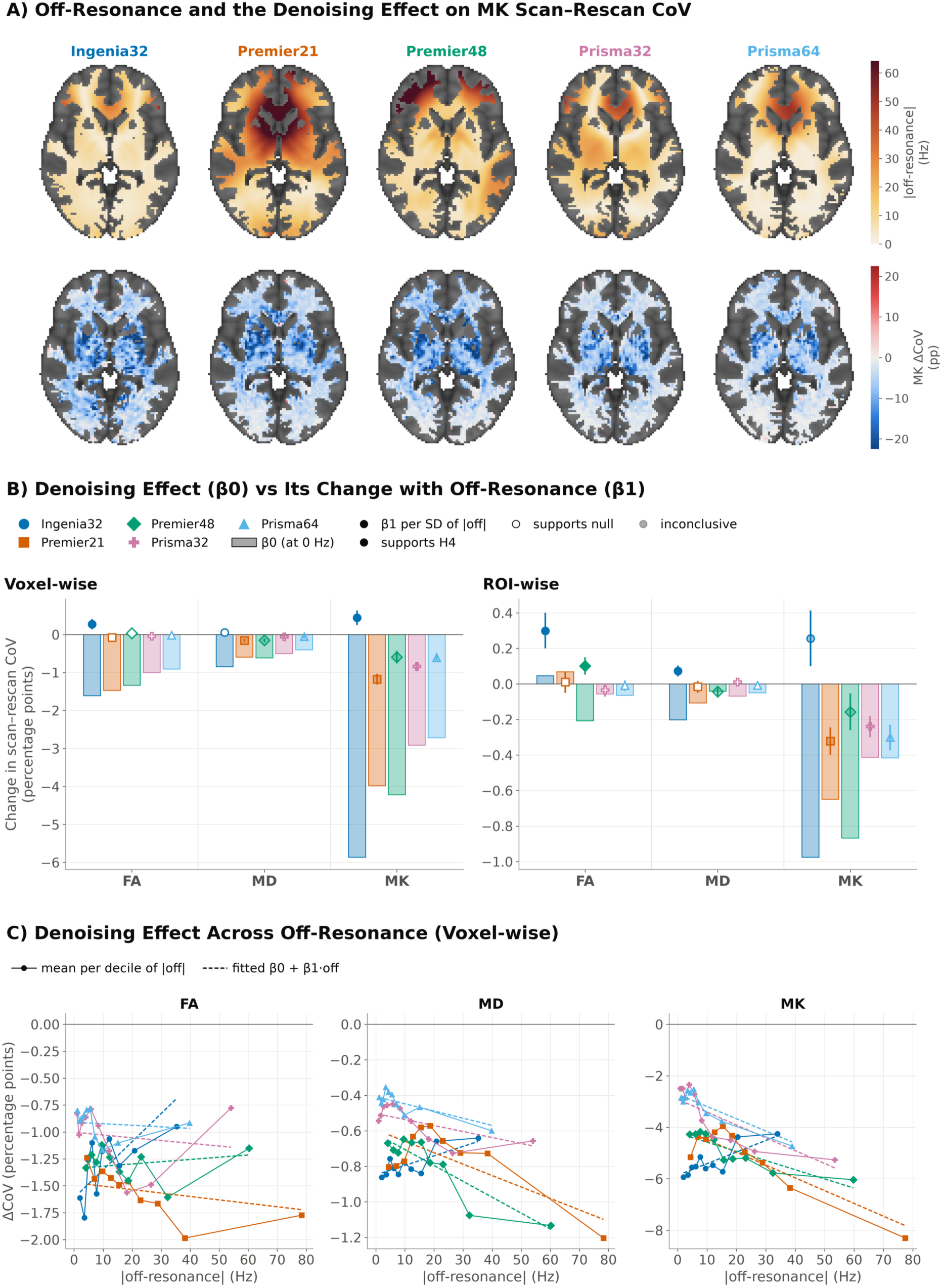
H4: Off-resonance and the effect of denoising on scan–rescan variability. (A) Mean absolute off-resonance (TOPUP field) over the six repeats (top) and ΔCoV = denoised − non-denoised scan-rescan CoV of MK (bottom). White-matter voxels warped to MNI space for display (the analysis used the checkerboard voxels in each subject’s template space). (B) The pre-registered regression. Bars: β0, the denoising effect at zero off-resonance. Markers: β1, the change in that effect per SD of |off-resonance|. (C) Voxel-wise ΔCoV against off-resonance, with the fitted line (dashed).

### 4.2 Exploratory Analysis

#### H1: Cross-scanner CoV changes with denoising

To understand why ROI-wise FA CoV increased, we split each subject’s mean CoV² (worked example, Figure 6A), into the cross-scanner variance of the mean across units (voxels/ROIs) and the within-scanner variance of units around those offsets (Figure 6B, Supplementary Table S14). The ROI-wise FA increase was present in all 10 subjects (median ΔCoV +0.16 to +0.45; 61–72% of ROIs). It came mainly from scanner offsets, which accounted for most of the rise in FA CoV², with little change in within-scanner variance. Voxel-wise, offsets increased similarly, but within-scanner variance decreased, resulting in a lower total CoV².

**Figure 6.**
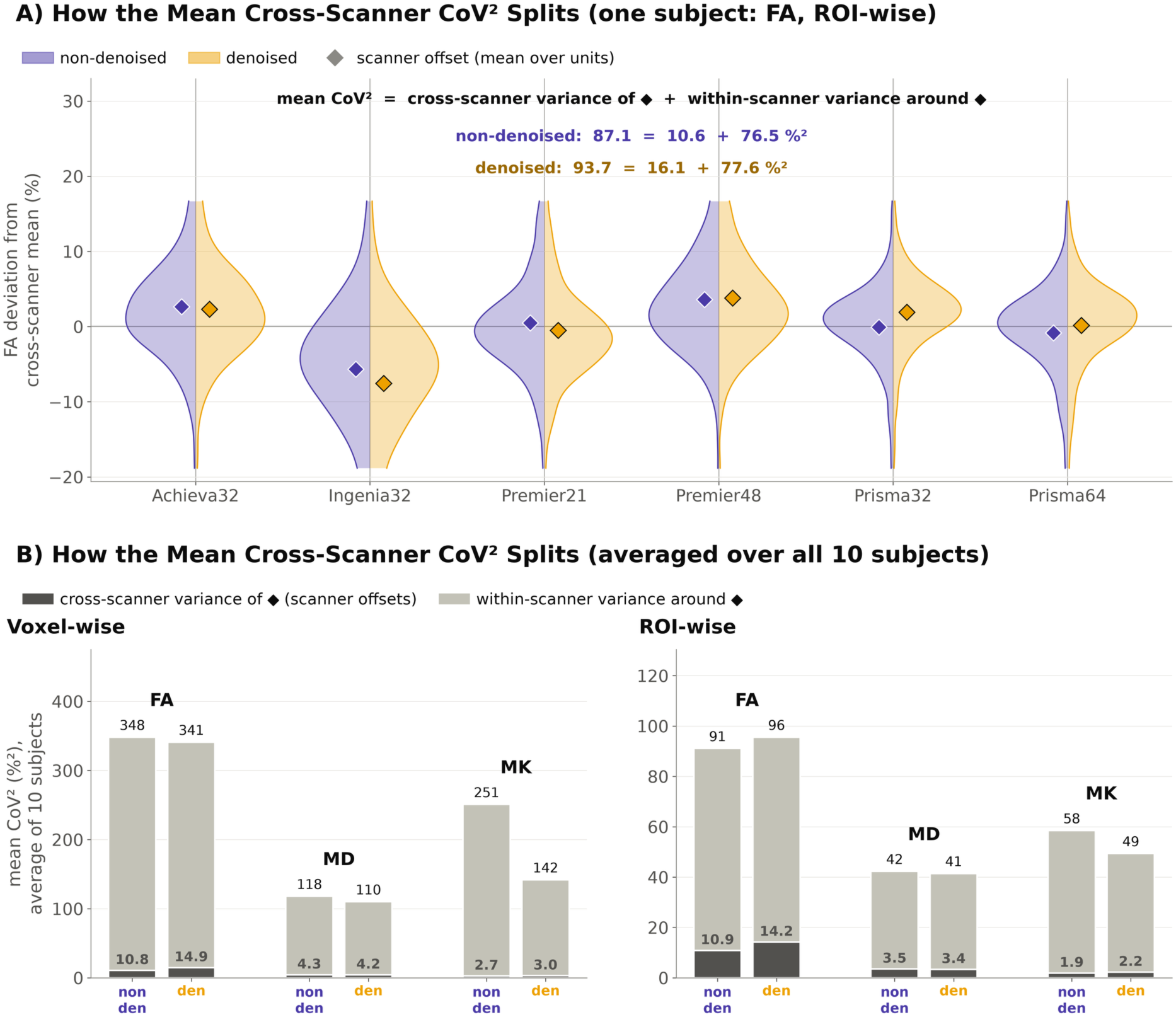
Cross-scanner CoV changes with denoising (exploratory). (A) Worked example for one subject. The per-unit CoV², averaged over units, splits exactly into the cross-scanner variance of the mean and the within-scanner variance of the units around their mean. (B) The same split for every metric and level, averaged over the 10 subjects. Dark = cross-scanner variance of the mean, light = within-scanner variance around the mean.

## H2: Cross-Scanner Bias

There was a strong relationship between the change in bias (Figure 3) and scanner CNR (Figure 7). Across subject and scanner combinations, denoising lowered FA and MD and raised MK, and the ROI-wise shift was larger at lower CNR (Figure 7A; Spearman ρ with CNR: FA +0.93, MD +0.91, MK −0.95). This relationship was also present within scanners, after removing scanner means (FA +0.84, MD +0.68, MK −0.78). Because bias compares the repeat scanner with the subject’s acquisitions from other scanners, a scanner with lower CNR than the others shifts further than others. Accordingly, ΔBias varied with the repeat scanner’s relative CNR in the five pairs (Figure 7B): pairs with relative CNR below 1 (GE, Philips) had negative FA and MD ΔBias and positive MK ΔBias, and the two Siemens pairs (above 1) had the reverse. Re-run with four times as many draws, three of the seven H2b fits that had not converged met the criteria; α0 changed by at most 0.004 and all verdicts were unchanged (Supplementary Table S15a).

**Figure 7.**
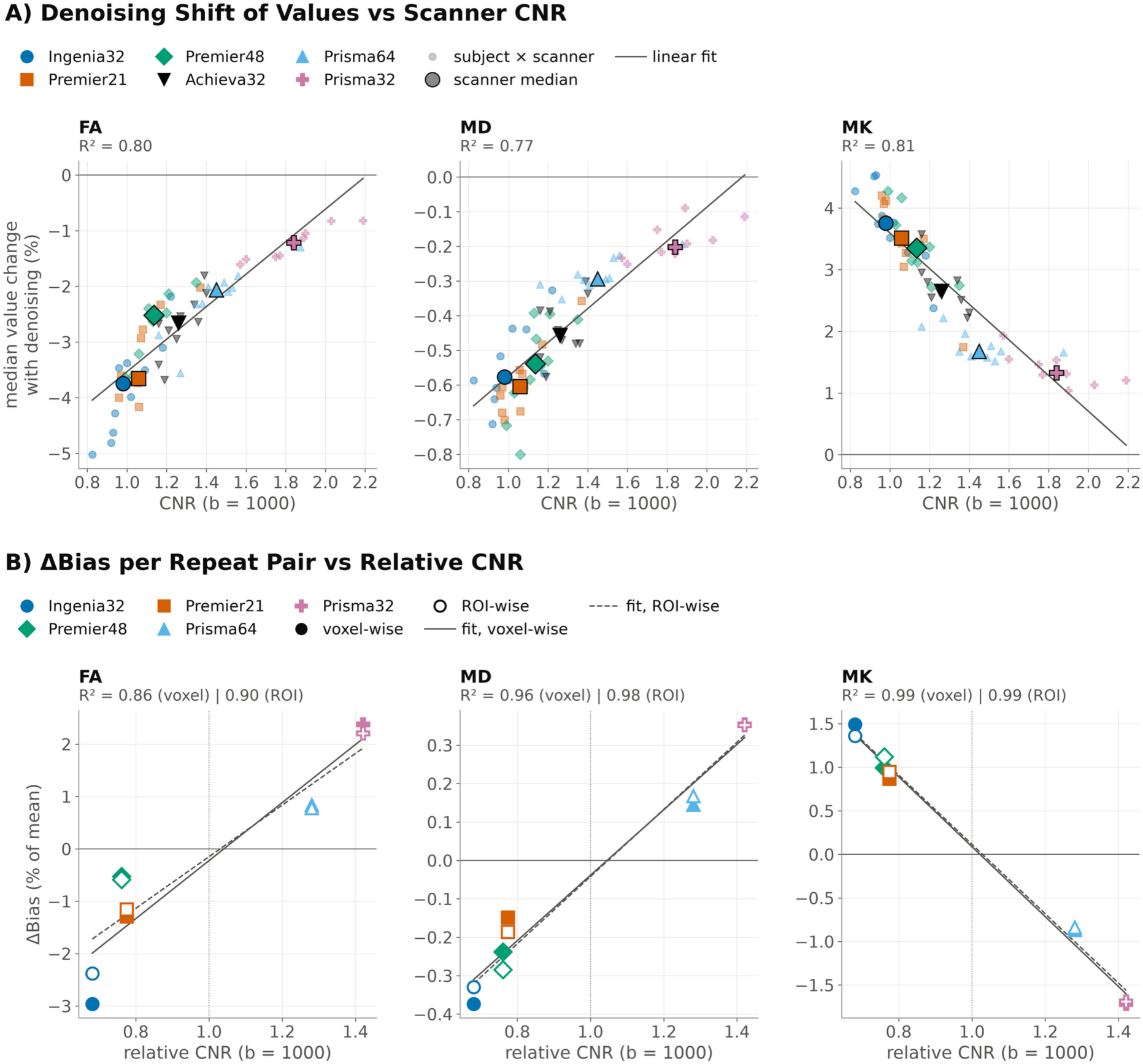
Relationship between the effect of denoising on metric values and scanner CNR (exploratory). (A) ROI-wise denoising shift per subject and scanner vs CNR (b=1000; EDDY QC, non-denoised), with the least-squares fit and its R². (B) ΔBias per repeat pair vs relative CNR. Relative CNR = repeat scanner (6 repeat sessions) ÷ mean of the subject’s other 5 scanners. Filled = voxel-wise, open = ROI-wise, lines = least-squares fits (solid = voxel-wise, dashed = ROI-wise), with R² under each metric.

## H3: Cross-Scanner Reliability

The registered model uses one subject and one scanner effect for all units, so differences between voxels/ROIs due to spatially varying anatomy fall into its residual (79–96% of variance). It therefore cannot detect changes in reliability at individual voxels or ROIs. We computed ICC(2,1) separately for each unit (Figure 8A). Per-unit ICC was much higher than the registered ICC (Figure 8B), and denoising further increased it, most strongly for voxel-wise MK (median per-unit ΔICC of +0.185, higher in 92% of voxels). Increases were small for voxel-wise FA (+0.019) and MD (+0.008), and for ROI-wise FA (+0.006) and MK (+0.043). The gain came from a smaller residual share (voxel-wise MK median 0.67 to 0.43; Figure 8B), while its scanner share did not fall (0.02 to 0.05). For voxel-wise MK, this used the 27% of voxels valid in all sessions (Supplementary Table S7). Re-run with four times as many draws, the two H3 tests that had not converged (voxel-wise MD and MK) did so and remained inconclusive (Supplementary Table S15b).

**Figure 8.**
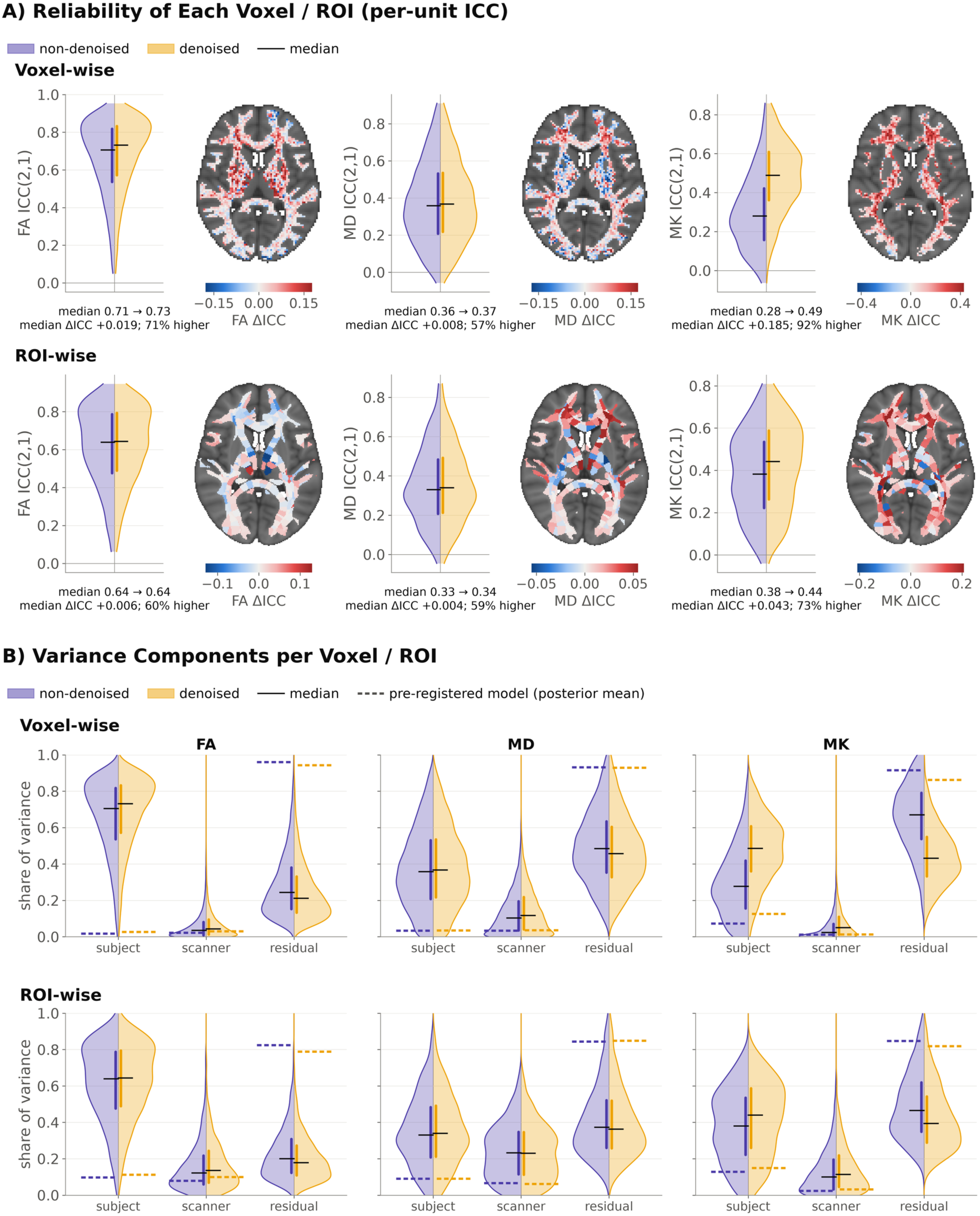
Reliability of each voxel and ROI (exploratory). (A) ICC(2,1) computed separately for each voxel/ROI from the 10 subjects × 6 scanners. Black line = median, bar = interquartile range. The violins and numbers use the checkerboard voxels valid across all 85 sessions. For display, the voxel-wise map shows all valid white-matter voxels. (B) Share of each unit’s variance due to subjects, scanners and the residual (two-way random-effects ANOVA per voxel/ROI), non-denoised (left half) and denoised (right half). The means of the pre-registered model are the dashed lines.

## H4: Denoising Association with Off-Resonance Fieldmaps

The registered models treated voxels and ROIs as independent. However, the off-resonance field varies smoothly across the brain, and spatial autocorrelation inflates the apparent significance of correlations between brain maps (Burt et al., 2020), so the credible intervals and Bayes factors overstate the precision of β1. To estimate how large a chance association such maps could produce, we generated 1,000 surrogate off-resonance fields (Figure 9A) for each repeat pair, each with the same values as the real field and a matched variogram (BrainSMASH; Burt et al., 2020), and recomputed the correlation between ΔCoV and |off-resonance| (Spearman ρ) for every surrogate. The surrogates reproduced the autocorrelation of the real field at distances beyond about 10mm and were slightly less smooth at shorter distances (Figure 9B).

**Figure 9.**
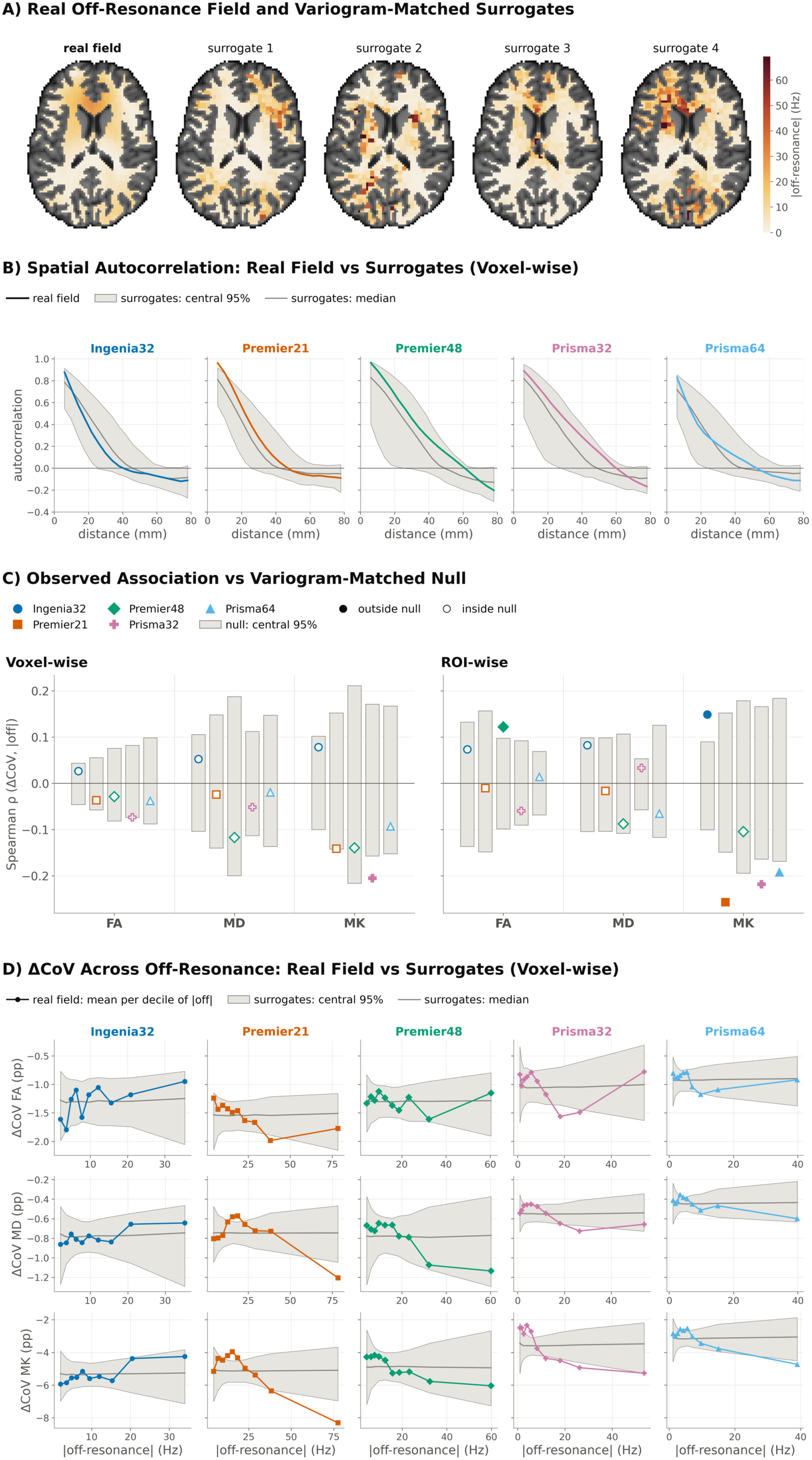
Variogram-matched null for the off-resonance association (exploratory). (A) Mean |off-resonance| of one repeat pair (Prisma64) on its white-matter voxels, and four surrogate fields with the same values and a matched variogram (BrainSMASH; Burt et al., 2020). (B) Spatial autocorrelation of the rank-transformed field against distance, voxel-wise, for each repeat pair: real field (coloured) and surrogates (grey line = median, band = central 95%). (C) Correlation (Spearman ρ) of ΔCoV with |off-resonance|: marker = observed ρ, grey bar = central 95% of ρ over 1,000 surrogate fields. (D) Voxel-wise ΔCoV against |off-resonance|, as in Figure 5C, with the central 95% and median over the surrogate fields.

The central 95% of chance correlations spanned ±0.04 to ±0.22 (Figure 9C), depending on the pair, metric and level. With 30 tests (five subject-scanner pairs x two levels, voxel- and ROI-wise x three metrics), one or two associations would be expected to fall outside this range by chance alone, six did. Two of these were in the direction predicted by H4, and both were weak: ROI-wise MK for the Ingenia32 pair (ρ = 0.15) and ROI-wise FA for the Premier48 pair (ρ = 0.12). The other four were in the opposite direction (ρ = −0.19 to −0.26), all of them for MK. Tested against the same surrogates (Supplementary Section 8), the registered slope β1 gave a similar picture (Supplementary Figure S16). Z-scoring |off-resonance| changed one H4 verdict, from null to inconclusive (Supplementary Table S17). These analyses therefore give no consistent evidence that off-resonance limits the benefit of denoising.

## 5. Discussion

Across scanners, MPPCA denoising lowered voxel-wise coefficients of variation for all three metrics, most strongly for kurtosis. The registered test of reliability was inconclusive, and, contrary to our hypothesis, denoising changed cross-scanner bias in every pair. Exploratory analyses suggest that denoising reduced the random component of cross-scanner variability and increased the reliability of individual voxels and ROIs, but it did not consistently reduce systematic differences between scanners, and that the size of the shift in metric values was strongly related with each scanner’s CNR. Registered and exploratory analyses together gave no convincing evidence that susceptibility-induced off-resonance limits the benefit of denoising. Our results suggest that denoising is beneficial for reducing noise-driven variance in multi-vendor data, but not for removing systematic scanner differences.

The voxel-wise reductions in CoV in the registered analyses extend findings within a single vendor (Ades-Aron et al., 2025) to scanners from three vendors. The effects were smaller ROI-wise, suggesting that averaging within an ROI already removes much of the thermal noise. They were largest for MK, the metric most affected by noise (Veraart et al., 2011). The registered H3 analysis was inconclusive: its model shares one subject and one scanner effect across all units, so 79–96% of the variance fell into the residual. In exploratory analyses, where ICC was computed separately for each unit, reliability followed the same pattern as CoV: it increased with denoising, most clearly for voxel-wise MK, because the residual share fell while the scanner share did not. Denoising may therefore improve sensitivity to differences between individuals without reducing differences between scanners.

The registered ROI-wise FA result was in the opposite direction to our hypothesis, and exploratory analyses showed that CoV increased in all ten subjects. Most of this increase came from larger offsets between scanners, with little change in the variance around them. This matches the possibility that removing noise exposes hardware- and software-related differences between manufacturers more clearly, although the CNR-related shifts described below may also contribute. If denoising does expose such software- and hardware-related differences, it could, speculatively, help harmonisation methods to estimate and remove them, as scanner effects would be less obscured by noise.

At low SNR, thermal noise inflates FA and biases MD and MK (Jones & Basser, 2004; Pierpaoli & Basser, 1996; Veraart et al., 2011). Removing it would therefore be expected to change values most on scanners and in sessions with low CNR. Our exploratory analyses are consistent with this explanation of the registered H2 results, as the change in bias was strongly related to the repeat scanner’s CNR relative to the subject’s other scanners. With five pairs this cannot be separated from vendor. However, the relationship between CNR and the size of the shift was also present across subject × scanner combinations and within scanners, which suggests it is not only a vendor effect, although other scanner, sequence or reconstruction differences may contribute. Whether such a shift brings a scanner closer to the others or moves it further away depends on the direction of its existing offset.

We found no consistent evidence that off-resonance limits denoising. Warrington et al. (2023) suggested that denoising is less effective where susceptibility artefacts are severe. Our results do not support a general effect of this kind. Five of 30 registered tests supported H4, four of them from the Ingenia32 pair, while twelve showed evidence in the opposite direction, mostly for MK. However, the registered models treated voxels and ROIs as independent, whereas the off-resonance field varies smoothly across the brain, so the Bayes factors overstate the evidence. In exploratory analyses against surrogate fields with matched spatial autocorrelation, one of the five associations supporting H4 exceeded chance when tested as rank correlations and four when tested as the registered slope; the rank correlations in the predicted direction were all weak (ρ ≤ 0.15). More effects exceeded chance in the opposite direction, mostly for MK, and support in the predicted direction came mainly from one pair (Ingenia32). Taken together, the registered and exploratory results do not give consistent evidence that susceptibility-related off-resonance limits the benefit of denoising.

These results have practical consequences for multi-site studies. Denoising reduces noise-driven variance, which many harmonisation tools are not explicitly designed to address, but our results suggest that systematic scanner offsets largely remain. Because the shift associated with denoising was related to CNR, we suggest caution when pooling denoised and non-denoised data, or when comparing denoised data from groups or sites that differ in CNR, as these may be shifted by different amounts.

## 6. Conclusion

MPPCA denoising reduced cross-scanner variability in voxel-wise diffusion metrics across vendors, most strongly for kurtosis. It did not consistently reduce systematic differences between scanners: ROI-averaged FA variability increased, and cross-scanner bias changed. Exploratory analyses indicated that this change was strongly related to scanner CNR, and that denoising improved the reliability of individual voxels by reducing residual variance. There was no convincing evidence that susceptibility-related off-resonance limits its benefit. Denoising is therefore useful for reducing random variance in multi-vendor studies, but it should complement rather than replace harmonisation tools.

## Data and Code Availability Statement

The ON-Harmony data are openly available for download from OpenNeuro (https://openneuro.org/datasets/ds004712). The code used in this report can be found in the GitHub repository (https://github.com/SPMIC-UoN/harmonisation-with-denoising).

## CRediT Author Contributions

FD: Conceptualisation, Methodology, Formal analysis, Software, Writing - original draft. JPMP: Conceptualisation, Writing - review and editing. OM: Supervision, Methodology, Writing - review and editing. SNS, PSM, SW: Conceptualisation, Methodology, Supervision, Writing - original draft, Writing - review and editing, Funding acquisition.

## Acknowledgements

FD, JPMP, OM, SW and SNS acknowledge support by an ERC Consolidator Grant (101000969). FD is also supported by a GE Healthcare co-funded PhD studentship. We thank Josefina Weinerova for her help in the statistical analysis plan.

