## Supplementary Material for "Does Denoising Reduce Cross-Vendor Variability in Diffusion MRI? A Travelling Heads Evaluation"

#### 1. GE Partial Fourier Fraction Calculation

GE data were acquired using the 'Min' partial Fourier setting. This specifies that 16 lines of k-space are acquired over half the matrix size. From Warrington et al., (2025), the matrix dimensions are 104x104 in-plane, therefore to calculate the partial Fourier fraction we calculate the following:

$$PF\% = \frac{52 + 16 \times 2}{104} \times 100 \approx 80.769\%$$

Where the multiplicative factor of two comes from the in plane parallel imaging factor.

**2. Scripts and parameters to be used for data preprocessing**

**Table S1:** parameters used for data preprocessing

| **Script** | **Use** | **Optional Parameters** | **Additional Information** |
|---|---|---|---|
| struc_preproc.sh | Preprocessing of T1 data | --qc | Part of the BRC pipelines (v1.6.3). |
| dMRI_preproc.sh | Preprocessing of dMRI data (including susceptibility, motion and eddy current correction) | --qc | |
| antsMultivariateTemplateConstruction2.sh | Creation of subject specific templates | | Part of ANTs toolbox (v2.4.4). |
| designer | Denoising dMRI data | -denoise<br>-extent 5,5,5<br>-adaptive_patch | Part of the DESIGNER pipeline (v2.0.11), used through the singularity container. |
| flirt | Linear registration of dMRI data to subject-specific T1 then subject-specific template. | -interp spline | Part of FSL toolbox (v6.0.6.4). |
| dtifit | Calculation of FA, MD, and MK parametric maps | --kurt | |
| fast | White matter segmentation on T1 subject-specific templates | | |
| fnirt | Non-linear registration to MNI template | | |
| MMORF | Multi-modal non-linear registration to MNI templates. The following templates will be used:<br>FSL_HCP1065_FA_1mm.nii.gz<br>FSL_HCP1065_tensor_1mm.nii.gz<br>MNI152_T1_1mm_brain.nii.gz | --modalities T1w,FA,DTI_tensors | Part of FSL toolbox (v6.0.7.18). |

### 3. Assessing the effect of denoising on susceptibility, eddy current and motion correction

Six within-scanner repeats of the Siemens Trio from Phase A of the ON-Harmony have been used to assess if denoising has a meaningful impact on susceptibility, eddy current, or motion corrections. This analysis was conducted to determine whether these corrections need additional modelling considerations dependent on denoising condition.

Parallel processing of denoised and non-denoised dMRI repeats was performed, involving susceptibility correction using FSL's TOPUP (Andersson et al., 2003) and eddy current and motion correction using FSL's EDDY (Andersson & Sotiropoulos, 2016). MPPCA denoising was carried out as described in the main manuscript.

Fieldmap values for each session were extracted from TOPUP correction and masked using a brain mask. The following metrics were extracted for each session and denoising condition: mean absolute field value, median absolute field value, standard deviation of field value, and 95th percentile value of the absolute field. Violin plots of these metrics are shown in Figure S2.

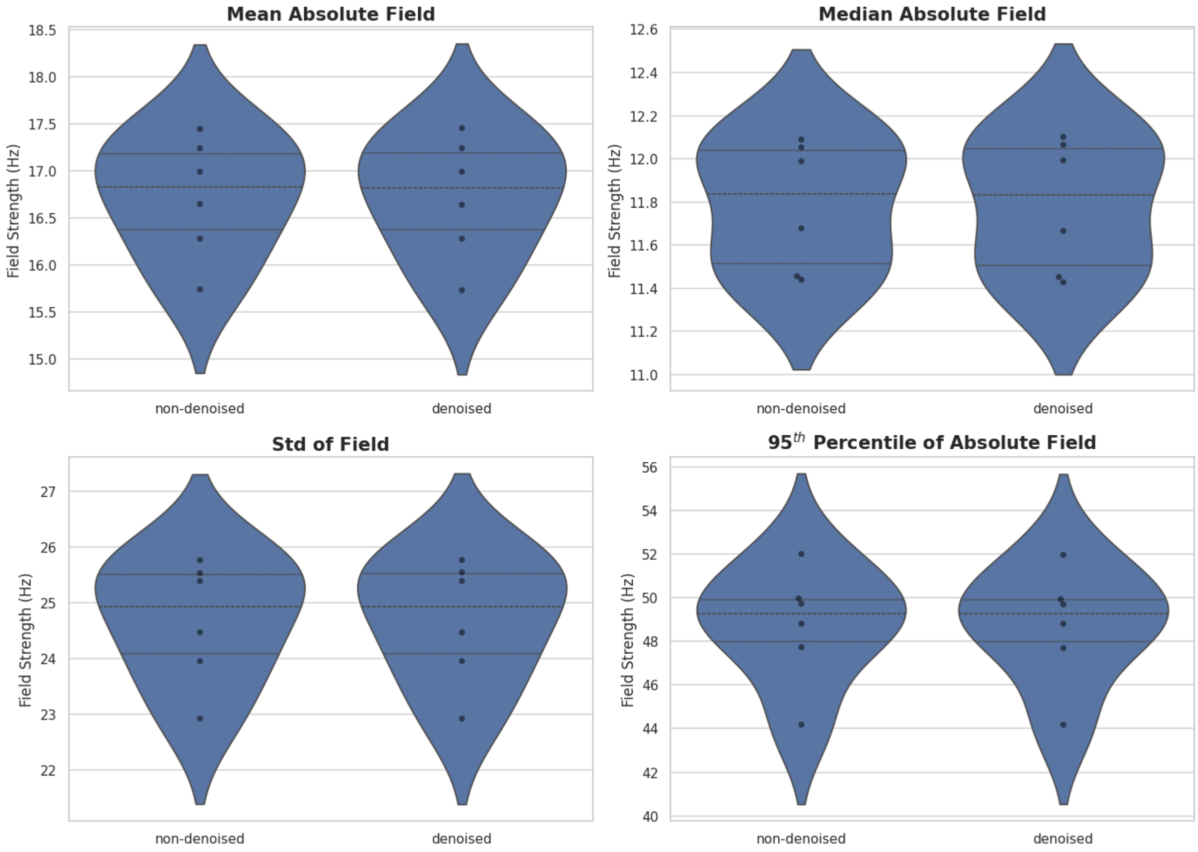


**Figure S2:** Visualisation of per-session susceptibility correction metrics.

To assess whether these values differed between denoising conditions, a Bayesian mixed-effects model was fitted separately for each of the four off-resonance metrics. The model was specified as:

$$metric \sim condition + (1|scan\_id)$$

where *metric* denotes the mean, median, standard deviation, or 95th percentile of the off-resonance field; condition indicates denoised versus non-denoised data; and (1 | scan_id) represents a random intercept for different scan sessions. Models were fit with 4 chains, 3000 iterations, and warmup = 1000; convergence was confirmed via R-hat and the absence of divergent transitions. The full code and outputs are available at https://github.com/SPMIC-UoN/harmonisation-with-denoising/tree/main.

Figure S3 shows the posterior distributions of the condition effect for each metric. In all cases, posterior means were close to zero and the 95% credible intervals spanned zero, indicating little evidence for a systematic difference between denoising conditions.

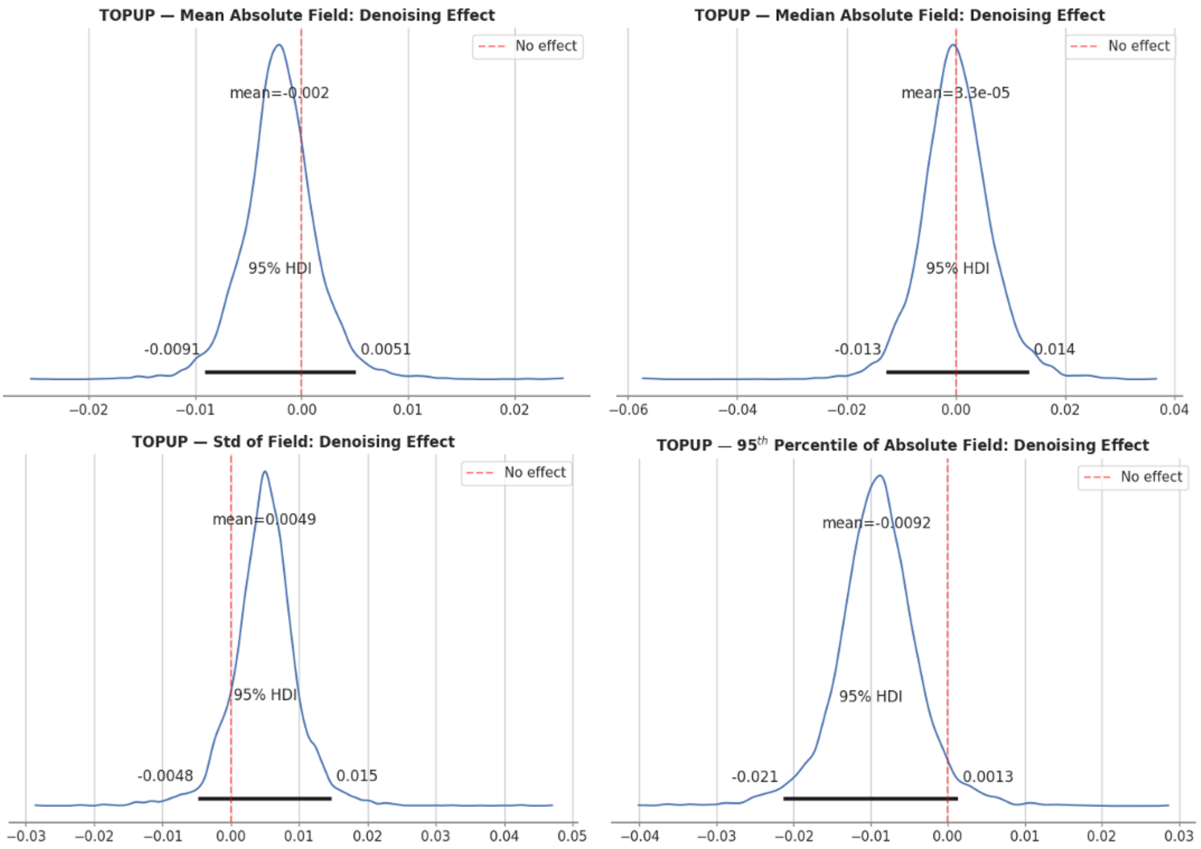


**Figure S3:** Posterior distributions of the effect of denoising on mean, median, standard deviation, and 95th percentile susceptibility fieldmap values within the brain. Posterior means were consistently near zero, with 95% credible intervals spanning zero, indicating minimal evidence for systematic differences between denoised and non-denoised conditions.

Eddy current and motion values for each session were extracted from EDDYQC (Bastiani et al., 2019) metrics, involving per-volume (109 volumes) measures of per-axis linear eddy current contributions and RMS movement. Violin plots of the pooled metrics from all six sessions are shown in Figure S4.

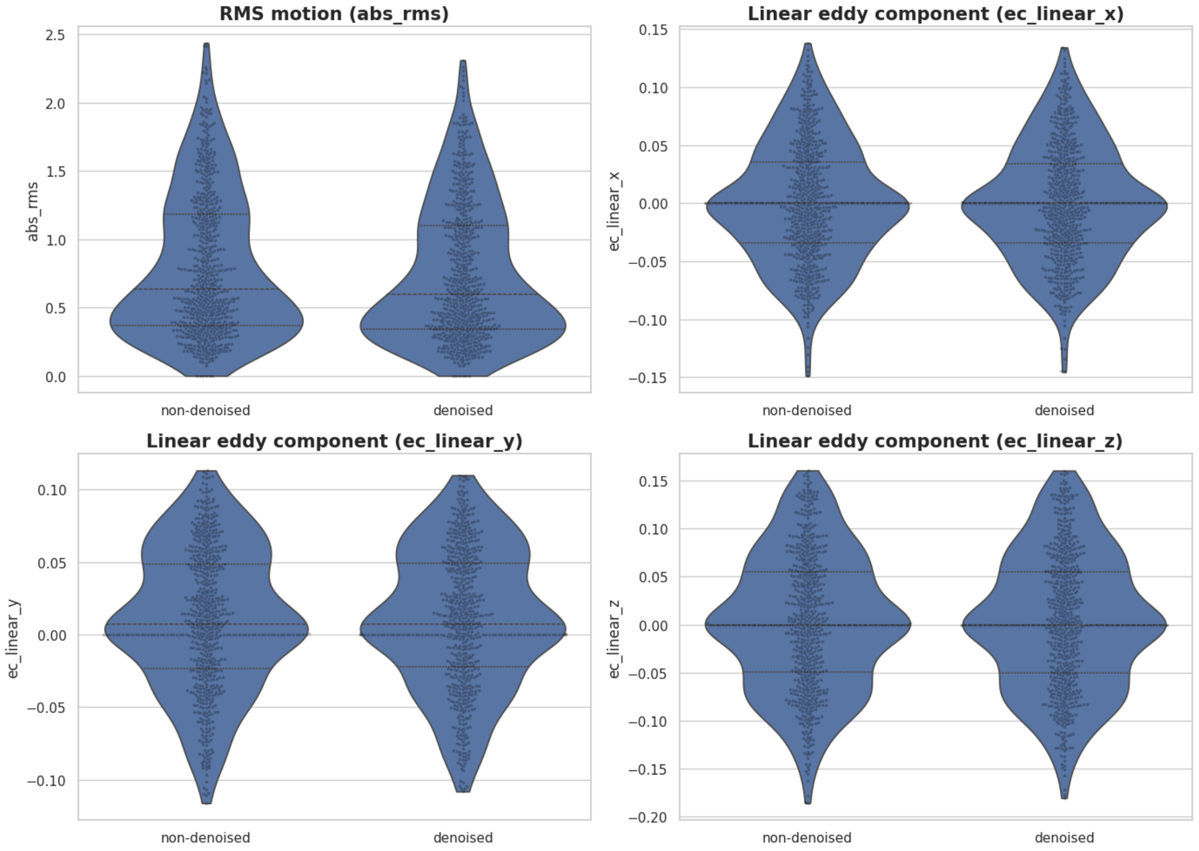


**Figure S4:** Visualisation of pooled eddy current and motion correction metrics across the six sessions.

The same Bayesian mixed-effects model ($metric \sim condition + (1|scan\_id)$) was applied to each of these EDDYQC metrics. Results are visualised in Figure S5, showing posterior means near zero with 95% credible intervals spanning zero, consistent with negligible effects of denoising.

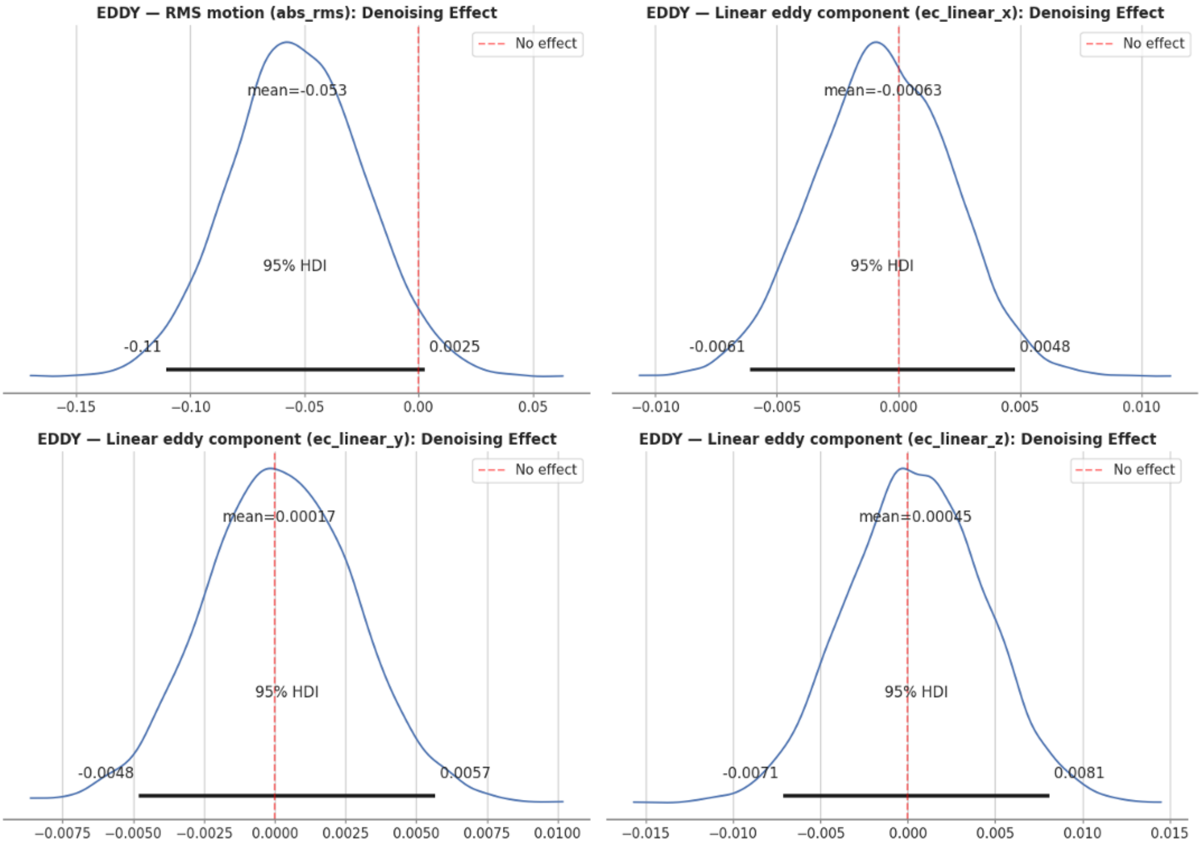


**Figure S5:** Posterior distributions of the effect of denoising on RMS motion and per-axis linear eddy currents. Posterior means are close to zero and 95% credible intervals span zero, indicating negligible differences between denoised and non-denoised data.

These results suggest that susceptibility, eddy current, and motion correction metrics are not affected by denoising and therefore do not require condition-specific consideration in models for H4.

### 4. Data Quality and Head Motion

**Table S6.** EDDY QC metrics per scanner (non-denoised data). Median [min–max] over sessions of mean absolute motion (relative to the first volume), mean relative motion (volume to volume), b = 0 SNR, and b = 1000 and b = 2000 s/mm² CNR. Achieva32 has 10 sessions (travelling heads only); the other scanners have 15 (10 travelling heads plus 5 additional repeats of one subject). For comparison, mean absolute motion was 0.70 ± 0.54 mm (median 0.56) in this study and 1.09 ± 0.47 mm (median 0.98) in UK Biobank (UK Biobank. Data-Field 24450: Mean absolute head motion from dMRI. UK Biobank Showcase. Retrieved 6 November 2026, from https://biobank.ndph.ox.ac.uk/showcase/field.cgi?id=24450).

| | Sessions | Absolute motion (mm) | Relative motion (mm) | SNR b0 | CNR b1000 | CNR b2000 |
|---|---|---|---|---|---|---|
| Achieva32 SPMICUP | 10 | 1.00 [0.44–4.53] | 0.28 [0.17–0.48] | 17.92 [16.77–20.98] | 1.26 [1.16–1.40] | 1.52 [1.28–1.64] |
| Ingenia32 SPMICUP | 15 | 0.57 [0.38–0.95] | 0.36 [0.24–0.80] | 15.46 [13.85–18.11] | 0.93 [0.78–1.22] | 0.97 [0.81–1.27] |
| Premier21 OCMR | 15 | 0.68 [0.31–1.47] | 0.25 [0.19–0.31] | 20.07 [17.53–21.56] | 1.07 [0.89–1.37] | 1.23 [0.97–1.69] |
| Premier48 SPMICQMC | 15 | 0.52 [0.34–0.78] | 0.22 [0.15–0.27] | 20.84 [16.12–23.34] | 1.13 [0.92–1.35] | 1.32 [1.08–1.51] |
| Prisma32 FMRIB | 15 | 0.40 [0.22–0.56] | 0.24 [0.22–0.30] | 24.59 [21.41–26.75] | 1.75 [1.45–2.19] | 1.87 [1.71–2.61] |
| Prisma64 OHBA | 15 | 0.61 [0.28–1.55] | 0.69 [0.20–0.80] | 23.35 [19.33–24.91] | 1.53 [1.16–1.92] | 1.90 [1.23–2.23] |
| All | 85 | 0.56 [0.22–4.53] | 0.26 [0.15–0.80] | 20.61 [13.85–26.75] | 1.20 [0.78–2.19] | 1.36 [0.81–2.61] |

### 5. Number of units kept

**Table S7.** Units per subject (H1), per subject–scanner pair (H2, H4) or common to all subjects (H3): mean (range) of the candidate units, and mean number kept (% of all candidate units, pooled over subjects or pairs). Voxels were excluded as unphysical, separately for each metric, where FA ≤ 0, MD ≤ 0, MK ≤ 0 or MK > 3 in either condition in any relevant session (main manuscript, Section 3.3). Voxel-wise models (a): one white matter voxel in eight; for H3a, voxels in MNI space valid in all 85 sessions. ROI-wise models (b): ROIs with at least one valid voxel (H3b: in all subjects).

| Model | Candidate units | FA (% kept) | MD (% kept) | MK (% kept) |
|---|---|---|---|---|
| H1a | 8,125 (6,301-9,397) | 7,781 (95.8%) | 7,841 (96.5%) | 7,230 (89.0%) |
| H1b | 1,298 | 1,257 (96.9%) | 1,259 (97.0%) | 1,253 (96.6%) |
| H2a | 8,376 (7,615-9,397) | 7,978 (95.2%) | 8,046 (96.1%) | 7,303 (87.2%) |
| H2b | 1,298 | 1,264 (97.4%) | 1,266 (97.5%) | 1,261 (97.1%) |
| H3a | 10,050 | 9,163 (91.2%) | 9,031 (89.9%) | 2,668 (26.5%) |
| H3b | 1,298 | 1,157 (89.1%) | 1,157 (89.1%) | 1,143 (88.1%) |
| H4a | 8,369 (7,624-9,271) | 8,294 (99.1%) | 8,294 (99.1%) | 7,966 (95.2%) |
| H4b | 1,298 | 1,275 (98.2%) | 1,275 (98.2%) | 1,273 (98.1%) |

### 6. Assessing the effect of denoising on susceptibility, eddy current and motion correction (Phase B Data)

The Stage 1 analysis (Section 3) was repeated on the full Phase B cohort: 85 sessions on 6 scanners. TOPUP and EDDY were run separately on the denoised and non-denoised data. The four field metrics were computed within the brain, using the non-denoised brain mask for both arms. The EDDY metrics were taken per volume (9,060 volumes per arm). Each metric was fitted with $metric \sim condition + (1|scan\_id)$ (bambi; TOPUP: 4 chains, 4,000 draws, 2,000 tuning steps; EDDY: 4 chains, 2,000 draws, 1,000 tuning steps).

Denoising did not meaningfully change the off-resonance field. The largest effect, on median |field|, was +0.016 Hz (95% HDI 0.003–0.029), 0.1% of its mean. The eddy-current terms did not change. EDDY's estimated RMS motion was 0.049 mm lower with denoising (95% HDI −0.059 to −0.040; 7%), well below the 2 mm voxel size.

The denoising effect converged for every metric (R-hat ≤ 1.01), but the session intercepts did not (R-hat up to 1.23); a paired comparison across sessions gave the same result (median |field| +0.016 Hz, 95% CI 0.003 to 0.029).

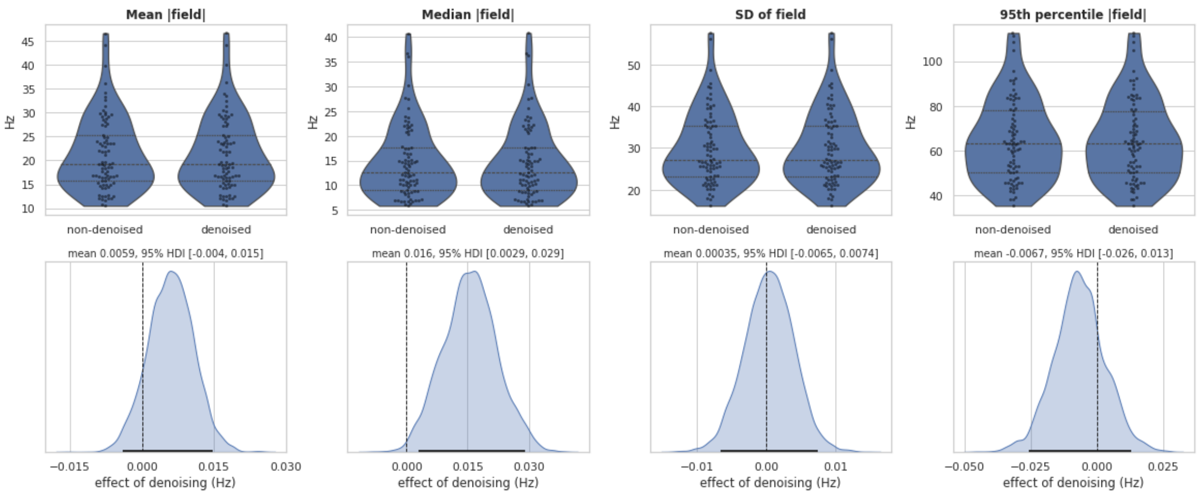


**Figure S8.** Effect of denoising on the TOPUP off-resonance field. Top: mean, median and 95th percentile of |field| and SD of the field within the brain, one point per session (85 sessions), for non-denoised and denoised data. Dashed lines inside the violins mark the median and quartiles. Bottom: posterior distribution of the denoising effect. The black bar is the 95% HDI and the dashed line is zero; the posterior mean and HDI are given above each panel.

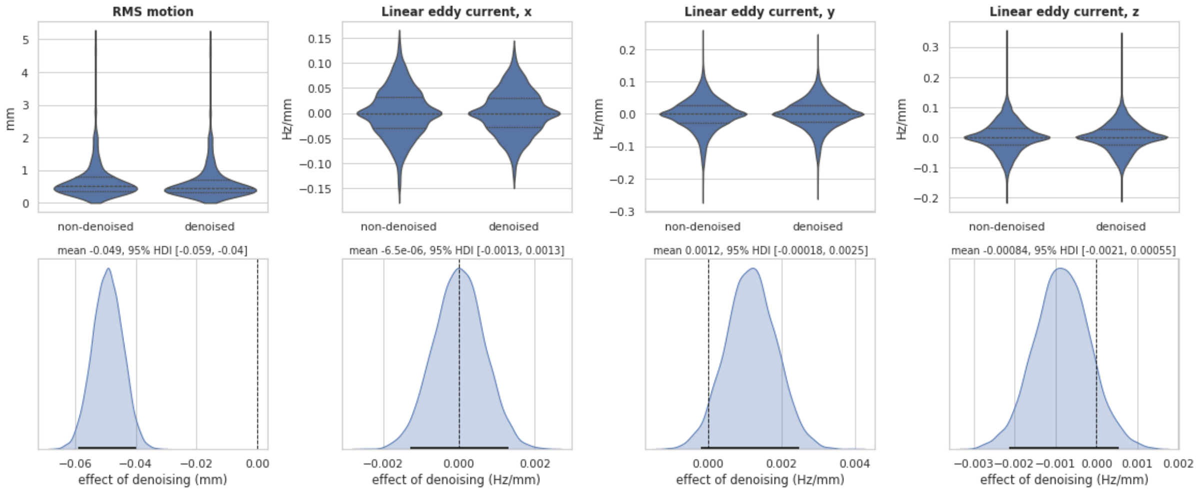


**Figure S9.** Effect of denoising on EDDY motion and eddy-current estimates. Top: absolute RMS motion (mm) and linear eddy-current terms along x, y and z (Hz/mm), per volume, pooled over the 85 sessions (9,060 volumes per arm). Bottom: posterior distribution of the denoising effect, as in Figure S8.

### 7. Registered Results

**Table S10.** H1: cross-scanner variability. α1: effect of denoising on z-scored CoV. Convergence: R-hat < 1.01 and effective sample size > 1,000, for the parameters used for inference.

| Hypothesis | α1 [95% CrI] | BF10 | median CoV (%), non-denoised → denoised | units with lower CoV (%) | converged | verdict |
|---|---|---|---|---|---|---|
| H1a – FA voxel-wise | −0.036 [−0.046, −0.026] | >100 | 13.0 → 12.6 | 56 | yes | supports H1 |
| H1a – MD voxel-wise | −0.074 [−0.084, −0.064] | >100 | 6.4 → 5.8 | 70 | yes | supports H1 |
| H1a – MK voxel-wise | −0.425 [−0.436, −0.415] | >100 | 10.0 → 6.8 | 81 | yes | supports H1 |
| H1b – FA ROI-wise | +0.060 [+0.044, +0.075] | >100 | 6.1 → 6.5 | 34 | yes | inconclusive (BF10 > 10, opposite sign) |
| H1b – MD ROI-wise | −0.023 [−0.040, −0.005] | 0.232 | 4.1 → 4.0 | 63 | yes | inconclusive |
| H1b – MK ROI-wise | −0.123 [−0.141, −0.106] | >100 | 4.6 → 4.1 | 66 | yes | supports H1 |

**Table S11.** H2: cross-scanner bias. One column per repeat pair (one subject scanned six times on that scanner). Cells: α0 [95% CrI] on ΔBias divided by its SD; mean ΔBias as % of the pair's mean value. BF10 > 100 for every test, so every verdict is "denoising changes bias". † did not meet the convergence criteria (re-run in Table S15a).

| α0 [95% CrI]; ΔBias (% of mean) | Ingenia32 | Premier48 | Premier21 | Prisma64 | Prisma32 |
|---|---|---|---|---|---|
| H2a – FA voxel-wise | −0.513 [−0.536, −0.491]; −3.0% | −0.161 [−0.184, −0.138]; −0.5% | −0.333 [−0.355, −0.311]; −1.3% | +0.287 [+0.264, +0.310]; +0.8% | +0.429 [+0.408, +0.450]; +2.4% |
| H2a – MD voxel-wise | −0.199 [−0.221, −0.176]; −0.4% | −0.147 [−0.169, −0.124]; −0.2% | −0.102 [−0.124, −0.080]; −0.1% | +0.122 [+0.101, +0.144]; +0.1% | +0.222 [+0.201, +0.243]; +0.4% |
| H2a – MK voxel-wise | +0.242 [+0.218, +0.265]; +1.5% | +0.185 [+0.161, +0.209]; +1.0% | +0.154 [+0.133, +0.177]; +0.9% | −0.168 [−0.191, −0.144]; −0.9% | −0.292 [−0.314, −0.270]; −1.7% |
| H2b – FA ROI-wise | −0.758 [−0.812, −0.702]; −2.4% | −0.409 [−0.459, −0.355] †; −0.6% | −0.625 [−0.679, −0.571] †; −1.2% | +0.636 [+0.582, +0.689] †; +0.8% | +1.230 [+1.176, +1.285]; +2.2% |
| H2b – MD ROI-wise | −0.396 [−0.453, −0.339] †; −0.3% | −0.491 [−0.545, −0.438]; −0.3% | −0.363 [−0.417, −0.312] †; −0.2% | +0.422 [+0.365, +0.479]; +0.2% | +0.685 [+0.628, +0.742]; +0.4% |
| H2b – MK ROI-wise | +0.540 [+0.486, +0.594]; +1.4% | +0.573 [+0.519, +0.629] †; +1.1% | +0.488 [+0.433, +0.543]; +0.9% | −0.455 [−0.513, −0.398] †; −0.8% | −0.840 [−0.895, −0.786]; −1.7% |

**Table S12.** H3: cross-scanner reliability. Test that did not meet the convergence criteria were re-run in Table S15b.

| Hypothesis | ICC non-denoised | ICC denoised | ΔICC [95% CrI] | P(ΔICC > 0) | converged | verdict |
|---|---|---|---|---|---|---|
| H3a – FA voxel-wise | 0.017 [0.006, 0.045] | 0.027 [0.010, 0.067] | +0.009 [−0.024, +0.052] | 0.747 | yes | inconclusive |
| H3a – MD voxel-wise | 0.034 [0.013, 0.084] | 0.034 [0.013, 0.088] | +0.000 [−0.055, +0.059] | 0.503 | no | inconclusive |
| H3a – MK voxel-wise | 0.073 [0.029, 0.163] | 0.126 [0.052, 0.278] | +0.052 [−0.064, +0.217] | 0.809 | no | inconclusive |
| H3b – FA ROI-wise | 0.097 [0.038, 0.224] | 0.112 [0.044, 0.252] | +0.015 [−0.131, +0.179] | 0.595 | yes | inconclusive |
| H3b – MD ROI-wise | 0.090 [0.036, 0.204] | 0.091 [0.036, 0.201] | +0.000 [−0.132, +0.132] | 0.498 | yes | inconclusive |
| H3b – MK ROI-wise | 0.129 [0.055, 0.268] | 0.149 [0.062, 0.309] | +0.021 [−0.151, +0.206] | 0.602 | yes | inconclusive |

**Table S13.** H4: off-resonance and the effect of denoising on scan–rescan CoV. Cells: β1 [95% CrI] (×$10^{-3}$ per Hz of |off-resonance|), BF10 and verdict. "H4": BF10 > 10 and β1 > 0; "null": BF10 < 1/10; "opposite": BF10 > 10 with β1 < 0 (inconclusive under the Stage 1 rules); "inconclusive": 1/10 < BF10 < 10. All fits converged.

| β1 [95% CrI] BF10 (verdict) | Ingenia32 | Premier48 | Premier21 | Prisma64 | Prisma32 |
|---|---|---|---|---|---|
| H4a – FA voxel-wise | +2.60 [+1.51, +3.72] BF 26.7 (H4) | +0.31 [−0.26, +0.91] BF <0.001 (null) | −0.51 [−1.02, −0.03] BF 0.002 (null) | −0.26 [−0.93, +0.40] BF <0.001 (null) | −0.35 [−0.94, +0.23] BF <0.001 (null) |
| H4a – MD voxel-wise | +0.96 [+0.37, +1.55] BF 0.048 (null) | −2.44 [−2.85, −2.01] BF >100 (opposite) | −1.52 [−1.75, −1.29] BF >100 (opposite) | −1.33 [−1.76, −0.91] BF >100 (opposite) | −0.99 [−1.33, −0.65] BF >100 (opposite) |
| H4a – MK voxel-wise | +4.76 [+2.93, +6.59] BF >100 (H4) | −4.43 [−5.50, −3.36] BF >100 (opposite) | −6.67 [−7.35, −5.98] BF >100 (opposite) | −6.86 [−7.96, −5.72] BF >100 (opposite) | −7.17 [−8.03, −6.32] BF >100 (opposite) |
| H4b – FA ROI-wise | +5.76 [+3.96, +7.64] BF >100 (H4) | +2.54 [+1.44, +3.64] BF 12.3 (H4) | +0.19 [−0.84, +1.21] BF <0.001 (null) | −0.33 [−1.20, +0.52] BF <0.001 (null) | −0.98 [−1.92, −0.03] BF 0.004 (null) |
| H4b – MD ROI-wise | +2.65 [+1.71, +3.60] BF >100 (H4) | −1.63 [−2.31, −0.97] BF 30.5 (opposite) | −0.22 [−0.62, +0.17] BF <0.001 (null) | −0.51 [−1.06, +0.03] BF 0.001 (null) | +0.61 [+0.09, +1.12] BF 0.004 (null) |
| H4b – MK ROI-wise | +5.15 [+2.10, +8.24] BF 0.377 (inconclusive) | −2.94 [−4.76, −1.08] BF 0.124 (inconclusive) | −5.07 [−6.23, −3.92] BF >100 (opposite) | −7.73 [−9.46, −6.01] BF >100 (opposite) | −6.32 [−7.78, −4.88] BF >100 (opposite) |

### 8. Exploratory Results

**Table S14. H1:** decomposition of the cross-scanner CoV². The mean squared CoV over units (%², averaged over the 10 subjects) splits exactly into the cross-scanner variance of the mean (the variance of the scanner offsets, i.e. each scanner's mean deviation from the cross-scanner mean) and the within-scanner variance of the units around those offsets. Shown for non-denoised and denoised data.

| metric | level | total denoised | total non-denoised | cross-scanner variance of the mean: denoised | cross-scanner variance of the mean: non-denoised | Within-scanner variance: denoised | Within-scanner variance: non-denoised |
|---|---|---|---|---|---|---|---|
| FA | ROI | 95.5 | 91.1 | 14.2 | 10.9 | 81.4 | 80.2 |
| FA | voxel | 340.9 | 348.1 | 14.9 | 10.8 | 326.0 | 337.3 |
| MD | ROI | 41.4 | 42.3 | 3.4 | 3.5 | 38.0 | 38.7 |
| MD | voxel | 110.0 | 117.9 | 4.2 | 4.3 | 105.8 | 113.6 |
| MK | ROI | 49.4 | 58.5 | 2.2 | 1.9 | 47.1 | 56.6 |
| MK | voxel | 141.8 | 250.8 | 3.0 | 2.7 | 138.8 | 248.1 |

**Table S15a.** H2b fits that did not meet the convergence criteria, with re-runs using four times as many draws (exploratory).

| Hypothesis | pair | draws | α0 [95% CrI] | BF10 | converged | verdict |
|---|---|---|---|---|---|---|
| H2b – FA ROI-wise | Premier48 | 8,000 per chain (4 chains) | −0.412 [−0.462, −0.356] | >100 | no | denoising changes bias |
| H2b – FA ROI-wise | Prisma64 | 8,000 per chain (4 chains) | +0.634 [+0.578, +0.689] | >100 | yes | denoising changes bias |
| H2b – FA ROI-wise | Premier21 | 8,000 per chain (4 chains) | −0.624 [−0.678, −0.571] | >100 | no | denoising changes bias |
| H2b – MD ROI-wise | Ingenia32 | 8,000 per chain (4 chains) | −0.392 [−0.449, −0.339] | >100 | no | denoising changes bias |
| H2b – MD ROI-wise | Premier21 | 8,000 per chain (4 chains) | −0.366 [−0.424, −0.316] | >100 | no | denoising changes bias |
| H2b – MK ROI-wise | Premier48 | 8,000 per chain (4 chains) | +0.575 [+0.521, +0.629] | >100 | yes | denoising changes bias |
| H2b – MK ROI-wise | Prisma64 | 8,000 per chain (4 chains) | −0.456 [−0.512, −0.401] | >100 | yes | denoising changes bias |

**Table S15b.** H3 fits that did not meet the convergence criteria, with re-runs using four times as many draws (exploratory).

| Hypothesis | draws | ICC non-denoised | ICC denoised | ΔICC [95% CrI] | P(ΔICC > 0) | converged | verdict |
|---|---|---|---|---|---|---|---|
| H3a – MD voxel-wise | 8,000 per chain (4 chains) | 0.034 [0.013, 0.083] | 0.034 [0.013, 0.086] | −0.000 [−0.055, +0.057] | 0.496 | yes | inconclusive |
| H3a – MK voxel-wise | 8,000 per chain (4 chains) | 0.072 [0.028, 0.169] | 0.127 [0.053, 0.280] | +0.055 [−0.066, +0.217] | 0.823 | yes | inconclusive |

**Surrogate Field Generation for H4**

Surrogate fields were generated with BrainSMASH v0.11 (Burt et al., 2020) from the rank-normalised |off-resonance| field. Voxel-wise, we used the 2,000 nearest sampled voxels of each voxel (median radius about 48 mm) and fitted the variogram over that whole range; ROI-wise, we used the distances between ROI centroids. These settings were chosen on one repeat pair, by how well the surrogates reproduced the autocorrelation of the real field, before the null distributions were computed. The same surrogates, with values mapped to those of the real field, were used for the null distribution of β1 (Figure S16).

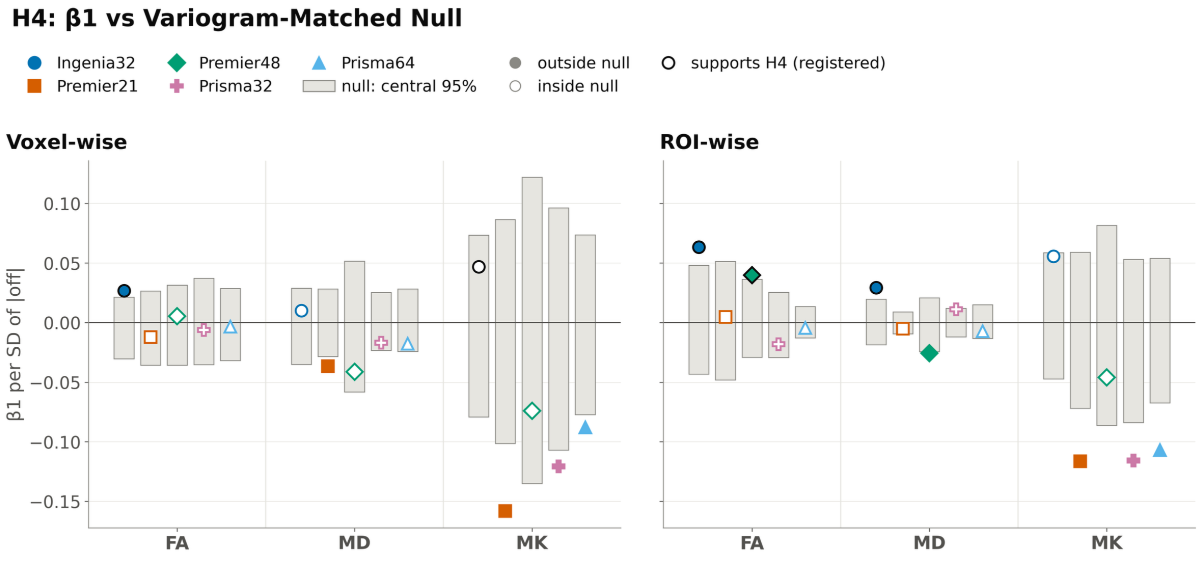


**Figure S16.** β1 against variogram-matched surrogate fields (H4, exploratory). β1 per SD of each pair's |off-resonance| (markers), which equals β1 of the model with z-scored |off-resonance|. Grey bars: central 95% of β1 over 1,000 surrogate fields with the same values and a matched variogram. Filled markers are outside that range (12 of 30: 4 with $\beta1 > 0$, as predicted by H4, and 8 with $\beta1 < 0$). Black outlines mark the five tests that support H4 in the registered analysis, four of which are outside the range.

**Table S17.** H4 verdict of each test with the main covariate (|off-resonance| in Hz) and with |off-resonance| z-scored within each pair (exploratory). Order within a cell: Hz / z-scored. Verdicts as in Table S13.

| Verdict: \|off\| Hz / \|off\| z | Ingenia32 | Premier48 | Premier21 | Prisma64 | Prisma32 |
|---|---|---|---|---|---|
| H4a – FA voxel-wise | H4 / H4 | null / null | null / null | null / null | null / null |
| H4a – MD voxel-wise | null / inconclusive | opposite / opposite | opposite / opposite | opposite / opposite | opposite / opposite |
| H4a – MK voxel-wise | H4 / H4 | opposite / opposite | opposite / opposite | opposite / opposite | opposite / opposite |
| H4b – FA ROI-wise | H4 / H4 | H4 / H4 | null / null | null / null | null / null |
| H4b – MD ROI-wise | H4 / H4 | opposite / opposite | null / null | null / null | null / null |
| H4b – MK ROI-wise | inconclusive / inconclusive | inconclusive / inconclusive | opposite / opposite | opposite / opposite | opposite / opposite |

### Dense Checkerboard Results

**Table S18.** Voxel-wise tests with one white matter voxel in eight (main analysis) and with the denser checkerboard (about 50% of white matter voxels; 25,246–37,527 per subject before exclusions). Estimates: α1, α0, ΔICC and β1 as in Tables S10–S13 (β1: $\times 10^{-3}$ per Hz). H1a, H2a, H4a: BF10; H3a: P(ΔICC > 0). † did not meet the convergence criteria. 35 of 36 verdicts are the same; the exception is H4a MD for Ingenia32.

| Test | Pair | 1 in 8: estimate [95% CrI] | 1 in 8: BF10 or P | 1 in 8: verdict | Dense: estimate [95% CrI] | Dense: BF10 or P | Dense: verdict |
|---|---|---|---|---|---|---|---|
| H1a FA (α1) | – | −0.036 [−0.046, −0.026] | >100 | supports H1 | −0.038 [−0.043, −0.033] | >100 | supports H1 |
| H1a MD (α1) | – | −0.074 [−0.084, −0.064] | >100 | supports H1 | −0.075 [−0.079, −0.070] | >100 | supports H1 |
| H1a MK (α1) | – | −0.425 [−0.436, −0.415] | >100 | supports H1 | −0.424 [−0.428, −0.419] | >100 | supports H1 |
| H2a FA (α0) | Ingenia32 | −0.513 [−0.536, −0.491] | >100 | denoising changes bias | −0.515 [−0.526, −0.503] | >100 | denoising changes bias |
| H2a FA (α0) | Premier48 | −0.161 [−0.184, −0.138] | >100 | denoising changes bias | −0.161 [−0.172, −0.150] | >100 | denoising changes bias |
| H2a FA (α0) | Premier21 | −0.333 [−0.355, −0.311] | >100 | denoising changes bias | −0.315 [−0.326, −0.304] | >100 | denoising changes bias |
| H2a FA (α0) | Prisma64 | +0.287 [+0.264, +0.310] | >100 | denoising changes bias | +0.292 [+0.281, +0.304] | >100 | denoising changes bias |
| H2a FA (α0) | Prisma32 | +0.429 [+0.408, +0.450] | >100 | denoising changes bias | +0.450 [+0.439, +0.460] | >100 | denoising changes bias |
| H2a MD (α0) | Ingenia32 | −0.199 [−0.221, −0.176] | >100 | denoising changes bias | −0.196 [−0.207, −0.185] | >100 | denoising changes bias |
| H2a MD (α0) | Premier48 | −0.147 [−0.169, −0.124] | >100 | denoising changes bias | −0.150 [−0.162, −0.139] | >100 | denoising changes bias |
| H2a MD (α0) | Premier21 | −0.102 [−0.124, −0.080] | >100 | denoising changes bias | −0.127 [−0.138, −0.116] | >100 | denoising changes bias |
| H2a MD (α0) | Prisma64 | +0.122 [+0.101, +0.144] | >100 | denoising changes bias | +0.136 [+0.125, +0.147] | >100 | denoising changes bias |
| H2a MD (α0) | Prisma32 | +0.222 [+0.201, +0.243] | >100 | denoising changes bias | +0.223 [+0.213, +0.234] | >100 | denoising changes bias |
| H2a MK (α0) | Ingenia32 | +0.242 [+0.218, +0.265] | >100 | denoising changes bias | +0.236 [+0.224, +0.248] | >100 | denoising changes bias |
| H2a MK (α0) | Premier48 | +0.185 [+0.161, +0.209] | >100 | denoising changes bias | +0.190 [+0.178, +0.201] | >100 | denoising changes bias |
| H2a MK (α0) | Premier21 | +0.154 [+0.133, +0.177] | >100 | denoising changes bias | +0.155 [+0.144, +0.166] | >100 | denoising changes bias |
| H2a MK (α0) | Prisma64 | −0.168 [−0.191, −0.144] | >100 | denoising changes bias | −0.168 [−0.180, −0.157] | >100 | denoising changes bias |
| H2a MK (α0) | Prisma32 | −0.292 [−0.314, −0.270] | >100 | denoising changes bias | −0.290 [−0.301, −0.280] | >100 | denoising changes bias |
| H3a FA (ΔICC) | – | +0.009 [−0.024, +0.052] | 0.75 | inconclusive | +0.010 [−0.023, +0.051] | 0.75 | inconclusive |
| H3a MD (ΔICC) | – | +0.000 [−0.055, +0.059] † | 0.50 | inconclusive | −0.000 [−0.064, +0.061] † | 0.51 | inconclusive |

| Test | Pair | 1 in 8: estimate [95% CrI] | 1 in 8: BF10 or P | 1 in 8: verdict | Dense: estimate [95% CrI] | Dense: BF10 or P | Dense: verdict |
|---|---|---|---|---|---|---|---|
| H3a MK (ΔICC) | – | +0.052 [−0.064, +0.217] † | 0.81 | inconclusive | +0.057 [−0.062, +0.223] † | 0.84 | inconclusive |
| H4a FA (β1) | Ingenia32 | +2.60 [+1.51, +3.72] | 26.7 | H4 | +2.30 [+1.77, +2.84] | >100 | H4 |
| H4a FA (β1) | Premier48 | +0.31 [−0.26, +0.91] | <0.001 | null | +0.19 [−0.11, +0.50] | <0.001 | null |
| H4a FA (β1) | Premier21 | −0.51 [−1.02, −0.03] | 0.002 | null | −0.01 [−0.26, +0.24] | <0.001 | null |
| H4a FA (β1) | Prisma64 | −0.26 [−0.93, +0.40] | <0.001 | null | +0.06 [−0.28, +0.40] | <0.001 | null |
| H4a FA (β1) | Prisma32 | −0.35 [−0.94, +0.23] | <0.001 | null | −0.49 [−0.78, −0.20] | 0.032 | null |
| H4a MD (β1) | Ingenia32 | +0.96 [+0.37, +1.55] | 0.048 | null | +1.35 [+1.08, +1.64] | >100 | H4 |
| H4a MD (β1) | Premier48 | −2.44 [−2.85, −2.01] | >100 | opposite | −2.50 [−2.73, −2.28] | >100 | opposite |
| H4a MD (β1) | Premier21 | −1.52 [−1.75, −1.29] | >100 | opposite | −1.34 [−1.44, −1.24] | >100 | opposite |
| H4a MD (β1) | Prisma64 | −1.33 [−1.76, −0.91] | >100 | opposite | −1.11 [−1.31, −0.90] | >100 | opposite |
| H4a MD (β1) | Prisma32 | −0.99 [−1.33, −0.65] | >100 | opposite | −1.27 [−1.45, −1.09] | >100 | opposite |
| H4a MK (β1) | Ingenia32 | +4.76 [+2.93, +6.59] | >100 | H4 | +5.47 [+4.58, +6.38] | >100 | H4 |
| H4a MK (β1) | Premier48 | −4.43 [−5.50, −3.36] | >100 | opposite | −4.09 [−4.63, −3.56] | >100 | opposite |
| H4a MK (β1) | Premier21 | −6.67 [−7.35, −5.98] | >100 | opposite | −6.65 [−7.00, −6.29] | >100 | opposite |
| H4a MK (β1) | Prisma64 | −6.86 [−7.96, −5.72] | >100 | opposite | −6.89 [−7.45, −6.33] | >100 | opposite |
| H4a MK (β1) | Prisma32 | −7.17 [−8.03, −6.32] | >100 | opposite | −6.82 [−7.24, −6.40] | >100 | opposite |
